# Inhibition of the gut ceramidase Asah2 decelerates the vertebrate ageing rate

**DOI:** 10.64898/2026.04.30.721799

**Authors:** Ayami Takaochi, Kota Abe, Yuki Sugiura, Akane Kawaguchi, Shigehiro Kuraku, Hide-Nori Tanaka, Daisuke Motooka, Kaori Tanaka, Farzana Ferdousi, Masao Nagasaki, Yasuyuki Ohkawa, Tohru Ishitani

**Affiliations:** Department of Homeostatic Regulation, Research Institute for Microbial Diseases, The University of Osaka, Suita, Osaka 565-0871, Japan; Multi-Omics Platform, Center for Cancer Immunotherapy and Immunobiology, Kyoto University Graduate School of Medicine, Kyoto, Kyoto 606-8501, Japan; Human Biology Microbiome Quantum Research Center, (WPI-Bio2Q), Keio University School of Medicine, Tokyo, 160-8582, Japan; Molecular Life History Laboratory, National Institute of Genetics, Mishima, Japan; Department of Genetics, SOKENDAI (Graduate University for Advanced Studies), Mishima, Japan; Institute for Glyco-core Research (iGCORE), Gifu University, Gifu, Gifu 501-1193, Japan; Genome Information Research Center, Research Institute for Microbial Diseases, The University of Osaka, Suita, Osaka 565-0871, Japan; Center for Infectious Disease Education and Research, The University of Osaka, Suita, Osaka 565-0871, Japan; Division of Transcriptomics, Medical Institute of Bioregulation, Kyushu University, Fukuoka, Fukuoka 812-0054, Japan; Division of Biomedical Information Analysis, Graduate School of Medical Sciences, Kyushu University, Fukuoka, Fukuoka 812-0054, Japan; Division of Biomedical Information Analysis, Medical Research Center for High Depth Omics, Medical Institute of Bioregulation, Kyushu University, Fukuoka, Fukuoka 812-0054, Japan; Center for Genomic Medicine, Graduate School of Medicine, Kyoto University, Kyoto, Kyoto 606-8507, Japan; AMED-CREST, The University of Osaka, Suita, Osaka 565-0871, Japan

**Author notes:** These authors are the equally contributed first authors (Ayami Takaochi, Kota Abe). Corresponding author: Tohru Ishitani; E-mail address: mailto.

**Keywords:** aging rate, systemic aging, comparative genomics, longevity, ceramide, microbiota, *N. furzeri*

## Abstract

The pace of ageing varies markedly among vertebrate species and individuals. However, the mechanisms underlying these differences in ageing rates remain unclear. Here, we show that the activity of the gut ceramidase Asah2 determines species- and strain-specific rates of vertebrate ageing. Comparative genomic analyses of vertebrate species and strains with different ageing rates reveal an association between low Asah2 activity and increased lifespan. Using the ultra-short-lived killifish *Nothobranchius furzeri* as a model, we demonstrate that knockout of Asah2 (*asah2* KO) extends lifespan and attenuates systemic ageing phenotypes, including declines in locomotor activity, abnormal protein accumulation in the brain, and accumulation of senescent cells in the liver. *asah2* KO elevates levels of ceramide species with long-chain fatty acids in the intestine, and supplementation with these ceramide species suppresses ageing phenotypes and extends lifespan in wild-type fish. *asah2* KO and ceramide supplementation alter gut microbiota composition, and *asah2* KO-derived microbiota transplantation attenuates ageing phenotypes, suggesting that reduced Asah2 activity prevents ageing through intestinal ceramide-mediated modulation of the microbiota. Given the evolutionary conservation of the Asah2 gene and its age-dependent upregulation in fish and humans, Asah2 and ceramides may act as ageing accelerators and decelerators, respectively, across animal species.

## Introduction

Lifespan varies dramatically across animal species, ranging from the African turquoise killifish (*Nothobranchius furzeri*), which lives for only a few months^1,2^, to the Greenland sharks, which lives for more than four centuries^3^. Such lifespan diversity has been utilised to investigate mechanisms of longevity^4^. For example, comparative genomic analysis of the long-lived Brandt’s bat and other bat species suggests that loss-of-function mutations in the growth hormone receptor may contribute to extended lifespan^5^. Comparative transcriptomic analyses of mammals with different lifespans have revealed that genes related to DNA repair and chromosome organisation are positively correlated with maximum lifespan (MLS)^6,7^, whereas expression of genes involved in energy metabolism, such as growth hormone (GH)/insulin-like growth factor signalling and mitochondrial metabolism, is negatively correlated with MLS^6,8–10^. Comparative metabolomic analyses have also identified distinct triacylglycerol species and alterations in sphingolipid metabolism in long-lived species^11,12^. However, because experimental validation of factors identified through comparative omics approaches is time-consuming, most remain candidate regulators of ageing.

*N. furzeri* is a powerful vertebrate model for comparative ageing research and intervention studies. It exhibits ageing-related characteristics similar to those observed in humans, including physiological and cognitive decline, and has an ultra-short lifespan that enables ageing analyses within a short timeframe^1,2^. Another significant advantage of *N. furzeri* is the existence of several strains that exhibit up to two-fold differences in lifespan^13^. Interestingly, although these strains do not differ in growth or maturation rates, their ageing rates differ significantly. Therefore, comparative analyses of these strains enable identification of regulators of ageing rate. Although quantitative trait locus mapping studies have revealed complex genetic architectures underlying lifespan differences^1,14^, it remains unclear which genes contribute to differences in ageing rates among *N. furzeri* strains.

Emerging evidence suggests that the intestine and its microbiota play pivotal roles in regulating systemic ageing. For example, gut-specific telomerase expression counteracts systemic ageing in telomerase-deficient zebrafish^15^. Furthermore, transplantation of gut microbiota from young adults prolongs lifespan in aged *N. furzeri* and attenuates systemic inflammation in mice^16,17^. Moreover, intestinal metabolites (e.g. lithocholic acid, a secondary bile acid, and the diet-derived metabolite urolithin A) have been shown to restore muscle strength in rodents^18–20^. However, the specific intestinal metabolites and enzymes that contribute to systemic ageing in vertebrates remain unclear. Here, we show that lower enzymatic activity of the gut ceramidase Asah2 is correlated with increased lifespan and that Asah2 accelerates systemic ageing through ceramide-mediated regulation of the microbiota.

## Results

### Loss-of-function mutations in Asah2 are associated with longevity

To explore previously unidentified regulators of ageing rate, we sequenced the whole genomes of the *N. furzeri* short-lived GRZ and long-lived MZM (MZM-0410) strains, which have median lifespans of 3–4 and 6–8 months, respectively^13^. This latest chromosome-scale GRZ genome assembly (NCBI Genomes, GCA_043380555.1) provides improved reconstruction of the karyotype and genome size, as well as higher coverage of protein-coding genes. Using this assembly as the reference for genome-wide alignment of MZM long reads (see Methods) enabled more comprehensive detection of small-scale variants and increased the number of single-nucleotide variants (SNVs) identified between the two strains (Supplementary Table 1a,b). A total of 7,053 SNVs were predicted to cause loss of protein function and were distributed across 1,669 genes (Supplementary Table 1c), 83 of which overlapped with known ageing-related genes (Supplementary Table 1d). These 1,669 genes were enriched in pathways related to gene expression, cell signalling, adhesion, and metabolism (Supplementary Fig. 1a; Supplementary Table 1e). We then focused on SNVs that met the following criteria: they were predicted to cause loss of protein function, were located in genes expressed exclusively in specific tissues, and occurred in genes encoding metabolic enzymes. The reason for setting these criteria is that tissue-specific metabolic enzymes represent promising targets for ageing interventions. Six genes were met these criteria (Supplementary Table 2a–c). Among them, we focused on the *asah2* gene, which encodes a neutral ceramidase specifically expressed in intestinal epithelial cells because the intestine plays an important role in ageing regulation^15–17^. The enzymatic domain of Asah2 is exposed to the extracellular space, facing the intestinal lumen^21^ (Fig. 1a). In the long-lived *N. furzeri* strain, the predicted Asah2 protein lacks its C-terminal domain owing to a single-nucleotide deletion (Fig. 1b; Supplementary Fig. 1b), which is essential for enzymatic activity^21^. By analysing the genomes of 464 animal species, we found that several relatively long-lived species within their respective classes^22^, such as *Myotis brandtii* and *Pantherophis guttatus* (Supplementary Fig. 1c), lack essential amino acids or domains required for Asah2 enzymatic activity, whereas humans and mice possess intact Asah2 (Fig. 1b). These findings suggest an association between reduced Asah2 activity and longevity; therefore, we focused on Asah2.

**Fig. 1:**
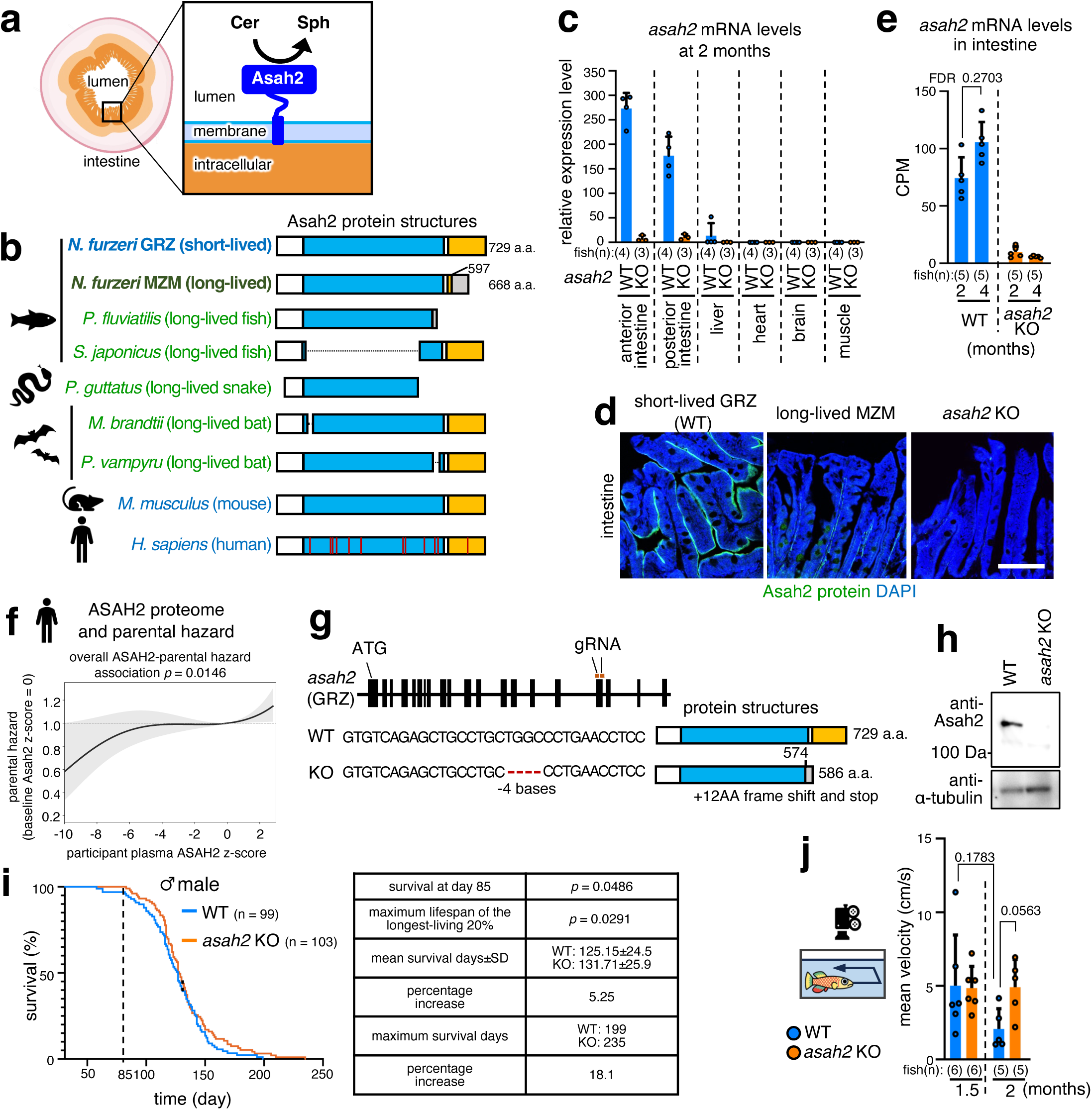
Loss of Asah2 activity extends vertebrate lifespan. **a**, Schematic representation of ceramide hydrolysis by membrane-tethered Asah2. Cer, ceramide; Sph, sphingosine. Image was created with BioRender.com. **b**, Loss of Asah2 activity is associated with a longer lifespan. Protein domain architectures of Asah2 are shown. Blue box, catalytic domain; orange box, immunoglobulin (Ig)-like domain; white box, other regions; grey box, *N. furzeri* MZM-specific region; red vertical lines in human ASAH2, essential amino acid residues required for enzymatic activity. **c, e**, *asah2* is specifically expressed in the intestine and increases with age. Expression levels of *asah2* mRNA in various tissues (**c**) and in the intestine (**e**) of 2- and 4-month-old wild-type (WT) and *asah2* knockout (KO) *N. furzeri* are shown. In **c** and **e**, mRNA levels were analysed by reverse transcription quantitative PCR (RT–qPCR) and RNA sequencing (RNA-seq), respectively. CPM, counts per million; FDR, false discovery rate. FDR values were derived from quasi-likelihood methods in edgeR. **d**, Asah2 protein localises to the apical /luminal side of intestinal epithelium in short-lived GRZ fish but is absent in long-lived MZM fish and *asah2* KO fish. Representative images of intestinal sections immunostained with anti-Asah2 antibody (green) and 4′,6-diamidino-2-phenylindole (DAPI; blue) are shown. Scale bar, 200 μm. **f**, Pooled censoring-aware parental survival analysis showing that higher participant plasma ASAH2 z-score was associated with higher pooled parental hazard, consistent with lower parental longevity. The displayed p-value tests the overall ASAH2-parental hazard association in the stratified cubic B-spline Cox model. Primary statistical inference was based on the cluster-robust linear Cox model. **g**, CRISPR/Cas9-mediated *asah2* knockout. Top, schematic representation of the *asah2* locus (black boxes, exons; ATG, start codon) and two sgRNA target sites (orange boxes). Bottom left, DNA sequences of WT and *asah2* KO. The red horizontal line denotes the 4 bp deletion. Bottom right, the mutation was predicted to result in loss of the C-terminal domain of the Asah2 protein. Blue, catalytic domain; orange, Ig-like domain; white, other regions; grey, *asah2* KO *N. furzeri-*specific region. **h**, Expression of Asah2 in the intestine of short-lived *N. furzeri* analysed by Western blotting. **i, j**, *asah2* KO extends lifespan (**i**) and improves locomotor activity (**j**). In **i**, survival curves of male WT (blue) and *asah2* KO (orange) fish are shown on the left; the corresponding statistical results, mean lifespan and maximum lifespan are shown on the right. Black dots indicate censored individuals. Graphs in **j** show exploratory behaviour in male WT (blue) and *asah2* KO (orange) fish. The y-axis indicates the average velocity. The log-rank test, fixed-time survival test, and unpaired two-tailed *t*-test were used for **i**, and a Brown–Forsythe and Welch ANOVA followed by Dunnett’s T3 multiple-comparisons test were used for **j**. Bars and error bars represent the mean ± SD.

### *asah2* KO extends male lifespan and healthspan

Consistent with observations in mice^23^, rats^24^, and humans^25–27^, *asah2* expression was detected specifically in the intestine of *N. furzeri* (Fig. 1c). Immunostaining revealed that, as in mice^23^ and humans^28^, Asah2 protein was localised to the apical/luminal side of the intestinal epithelium in the short-lived *N. furzeri* strain, whereas it was absent in the intestine of the long-lived strain (Fig. 1d). These findings suggest that the SNV in *asah2* in the long-lived strain leads to loss of intestinal Asah2 protein expression, rather than merely truncation of the C-terminal domain. Interestingly, *asah2* expression increased with age in male *N. furzeri* (Fig. 1e). Publicly available human data^29^ further showed that *asah2* expression in the terminal ileum of the small intestine increased with age (Supplementary Fig. 2a). Furthermore, in UK Biobank, higher participant plasma ASAH2 protein levels were associated with shorter parental survival in a pooled Cox analysis that included living parents as right-censored observations (overall 3-df spline association, p = 0.0146; cluster-robust linear Cox HR = 1.0183 per 1-s.d. higher ASAH2, 95% CI 1.0039–1.0329, p = 0.0125; Fig. 1f; Supplementary Fig. 2b). Given that plasma protein profiles are heritable and that human ASAH2 is specifically expressed in the intestine^25–27,30^, elevated ASAH2 levels may be associated with shorter lifespan in humans. These findings raise the possibility that Asah2 functions as an ageing accelerator contributing to lifespan differences among species and individuals. To test this hypothesis, we knocked out the *asah2* gene in the short-lived *N. furzeri* using the CRISPR/Cas9 system. The knockout (KO) fish harboured a four-base deletion in the *asah2* gene, resulting in a frameshift mutation and a premature stop codon upstream of the C-terminal region (Fig. 1g), which led to loss of Asah2 expression at both the mRNA and protein levels (Fig. 1c–e,h). The *asah2* KO did not affect body size, body mass index, or reproductive capacity (Supplementary Fig. 2c,d). The *asah2* KO extended lifespan and ameliorated the age-dependent decline in locomotor activity, which is considered an integrated measure of individual health^17^, in 2-month-old fish (Fig. 1i,j; Supplementary Movie 1). Thus, inhibition of Asah2 activity prevents organismal ageing.

### *asah2* KO prevents ageing in multiple organs

To further elucidate the effect of *asah2* KO on healthspan, we examined age-related phenotypes across tissues. As *asah2* KO prevented the age-dependent decline in locomotor activity, we first focused on the cerebellum and skeletal muscle, which regulate motor function. The *asah2* KO suppressed age-dependent accumulation of mono- and poly-ubiquitinated proteins, markers of impaired proteostasis^31^, in the cerebellum (Fig. 2a). The *asah2* KO also reduced the frequency of thin muscle fibres and increased the frequency of thick fibres in 2-month-old fish (Fig. 2b), suggesting that *asah2* KO ameliorates muscle ageing. In the brain, RNA sequencing (RNA-seq) followed by KEGG pathway and Gene Ontology (GO) term enrichment analyses showed that age-dependent upregulation of translation-related pathways and ribosomal protein genes was suppressed by *asah2* KO (Fig. 2c; Supplementary Fig. 3a,b). Similar age-dependent upregulation of ribosomal protein expression has been observed in the brains of mice, humans, and *N. furzeri*^32–34^. Indeed, an age-associated increase in ribosomal protein aggregation has been reported in the brains of mice and *N. furzeri*^34^, as well as in *Caenorhabditis elegans*^35^, and aberrant increases in ribosomal proteins abrogate proteostasis^36^. Thus, *asah2* KO may mitigate age-dependent decline in proteostasis by preventing abnormal increases in ribosomal proteins. We also found that, in the liver of 4-month-old fish, *asah2* KO blocked upregulation of ribosomal protein genes (Fig. 2c), which has recently been reported to be associated with a short-lifespan trajectory^37^, as well as accumulation of Senescence-associated β-galactosidase (SA-β-gal)-positive senescent cells (Fig. 2d), a hallmark of aging^31^.

**Fig. 2:**
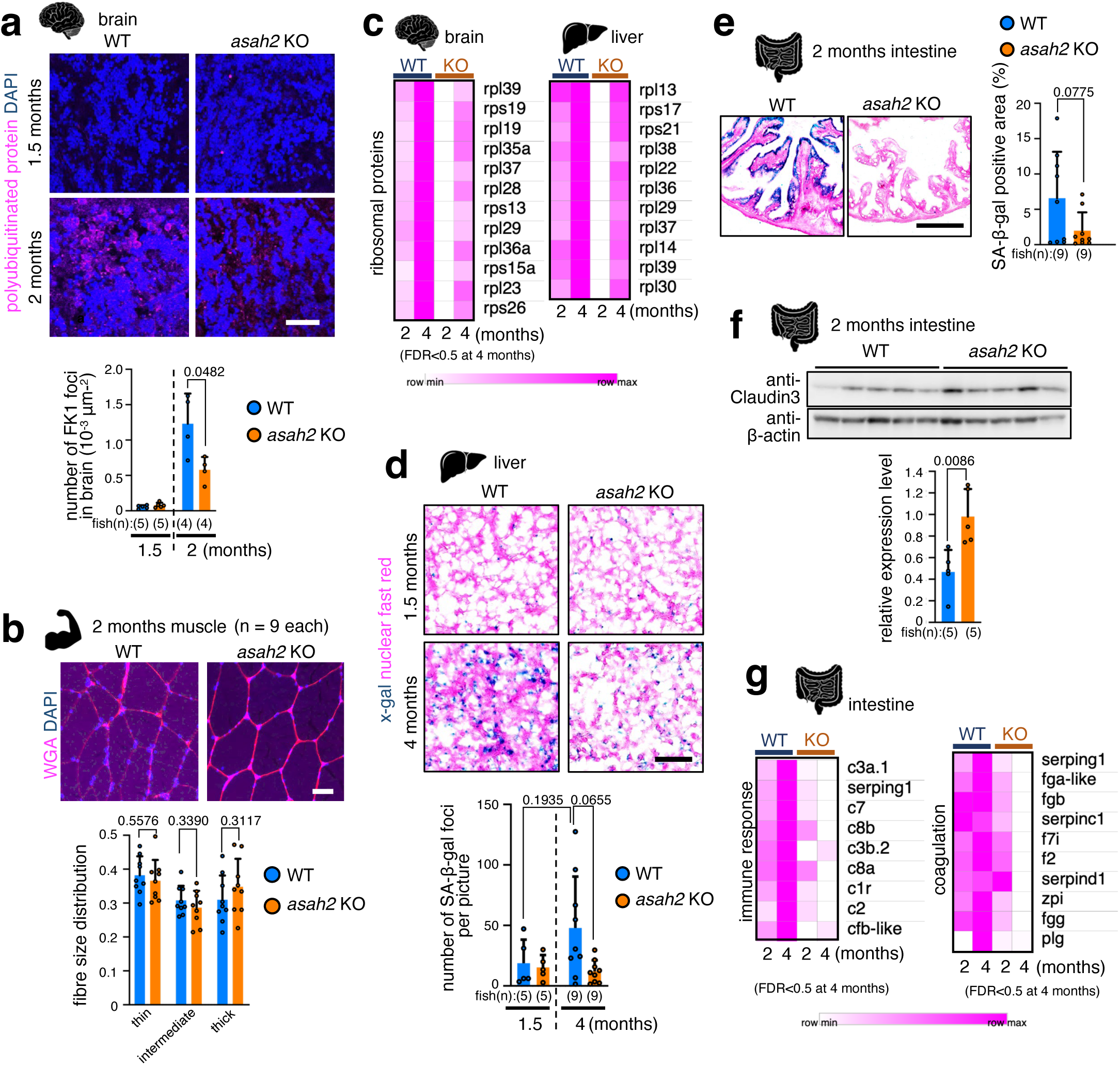
*asah2* KO prevents ageing in various tissues. **a**, *asah2* KO blocks age-dependent accumulation of polyubiquitinated proteins in the cerebellum. Representative images show cerebellar sections stained with the FK1 antibody (magenta) and 4′,6-diamidino-2-phenylindole (DAPI; blue) in 1.5- and 2-month-old fish. Scale bar, 50 μm. The graph shows the number of FK1 foci in WT (blue) and *asah2* KO (orange) fish. **b**, *asah2* KO blocks muscle fibre atrophy. Representative images show muscle sections stained with wheat germ agglutinin (WGA; red) and DAPI (blue) in 2-month-old fish. Scale bar, 50 μm. Graphs show the proportions of thin (< 6,000 μm^2^), intermediate (6,000–9,000 μm^2^), and thick (> 9,000 μm^2^) skeletal muscle fibres in WT (blue) and *asah2* KO (orange) fish. **c**, *asah2* KO blocks age-dependent upregulation of ribosomal protein genes in the brain and liver. mRNA levels were analysed by RNA sequencing (RNA-seq). **d**, *asah2* KO blocks age-dependent accumulation of senescent cells in the liver. Representative images show liver sections stained for senescence-associated β-galactosidase (SA-β-gal) and counterstained with Nuclear Fast Red. Scale bar, 50 μm. The graph shows the number of SA-β-gal-stained foci in the liver of WT (blue) and *asah2* KO (orange) fish. **e, f**, *asah2* KO blocks accumulation of senescent cells and increases Claudin-3 expression in the intestine of 2-month-old fish. In **e**, representative images show intestinal sections stained for SA-β-gal and counterstained with Nuclear Fast Red. The graph in **e** shows the percentage of SA-β-gal-positive area in WT (blue) and *asah2* KO (orange) fish. In **f**, Claudin-3 expression in the intestine of WT and *asah2* KO fish was analysed by Western blotting. The graph in **f** shows the relative expression of Claudin-3 in WT (blue) and *asah2* KO (orange) fish. **g**, *asah2* KO blocks age-dependent upregulation of immune response-and coagulation-related genes in the intestine. mRNA levels were analysed by RNA-seq. Bars and error bars (**a, b, d**–**f**) represent the mean ± SD. A Brown–Forsythe and Welch ANOVA followed by Dunnett’s T3 multiple-comparisons test was used in **a** and **d**, and an unpaired two-tailed *t*-test was used in **b, e,** and **f**. FDR values in **c** and **g** were derived from quasi-likelihood methods in edgeR. FDR, false discovery rate.

Next, we examined the intestine, where Asah2 is specifically expressed. We found that *asah2* KO reduced SA-β-gal-positive senescent cells in the intestine of 2-month-old fish (Fig. 2e). We also observed that *asah2* KO increased expression of Claudin-3, a major intestinal tight junction protein essential for barrier integrity^38^, in the intestine of 2-month-old fish (Fig. 2f); notably, Claudin-3 expression declines with age in the intestines of humans and mice^39^. It has been reported that reduced Claudin-3 expression leads to increased intestinal permeability (leaky gut) and consequent inflammation^38,40^. Thus, *asah2* KO may prevent age-dependent increases in intestinal permeability and inflammation. Consistent with this interpretation, GO term enrichment analysis of intestinal RNA-seq data showed that age-dependent upregulation of immune response- and blood coagulation-related genes was suppressed in *asah2* KO fish (Fig. 2g; Supplementary Fig. 3b), both of which are known to be activated by intestinal inflammation and are associated with chronic inflammatory disorders^41,42^. Since increased intestinal permeability and inflammation can promote systemic inflammation^16,43,44^, Asah2 may regulate ageing in multiple organs by inducing these intestinal alterations.

### *asah2* KO blocks ageing through increasing ceramide levels

As Asah2 is a neutral ceramidase that catalyses the hydrolysis of ceramides, we hypothesised that *asah2* KO prevents ageing by increasing ceramide levels. To test this hypothesis, we performed LC-HRMS-based lipidomic profiling of the intestine. Partial least squares discriminant analysis showed that *asah2* KO altered the intestinal lipidome (Fig. 3a), including levels of ceramides, sphingomyelins, and diacylglycerols (Supplementary Fig. 4a). As anticipated, *asah2* KO reduced the total amount of sphingosine, a metabolite of ceramide (Supplementary Fig. 4b). Although the total ceramide content did not change (Supplementary Fig. 4b), levels of several ceramide species, including Cer(d16:1/22:0) and Cer(d17:1/22:0), were substantially increased in the intestines of *asah2* KO fish (Fig. 3b). In line with the age-dependent increase in *asah2* levels, levels of Cer(d16:1/22:0) and Cer(d17:1/22:0) decreased with age (Supplementary Fig. 4c). To investigate the roles of Cer(d16:1/22:0) and Cer(d17:1/22:0) in ageing regulation, we synthesised these ceramides and administered them to 1.5-month-old fish via food (bloodworms) (Fig. 3c). Similar to *asah2* KO, ceramide supplementation extended lifespan (Fig. 3d) and prevented age-dependent decline in locomotor activity (Fig. 3e). Ceramide supplementation also reduced the frequency of thin muscle fibres, increased the frequency of thick fibres (Fig. 3f), and suppressed expression of immune response- and coagulation-related genes in the intestine of 4.5-month-old fish (Fig. 3g). These findings suggest that ceramides negatively regulate systemic ageing downstream of Asah2.

**Fig. 3:**
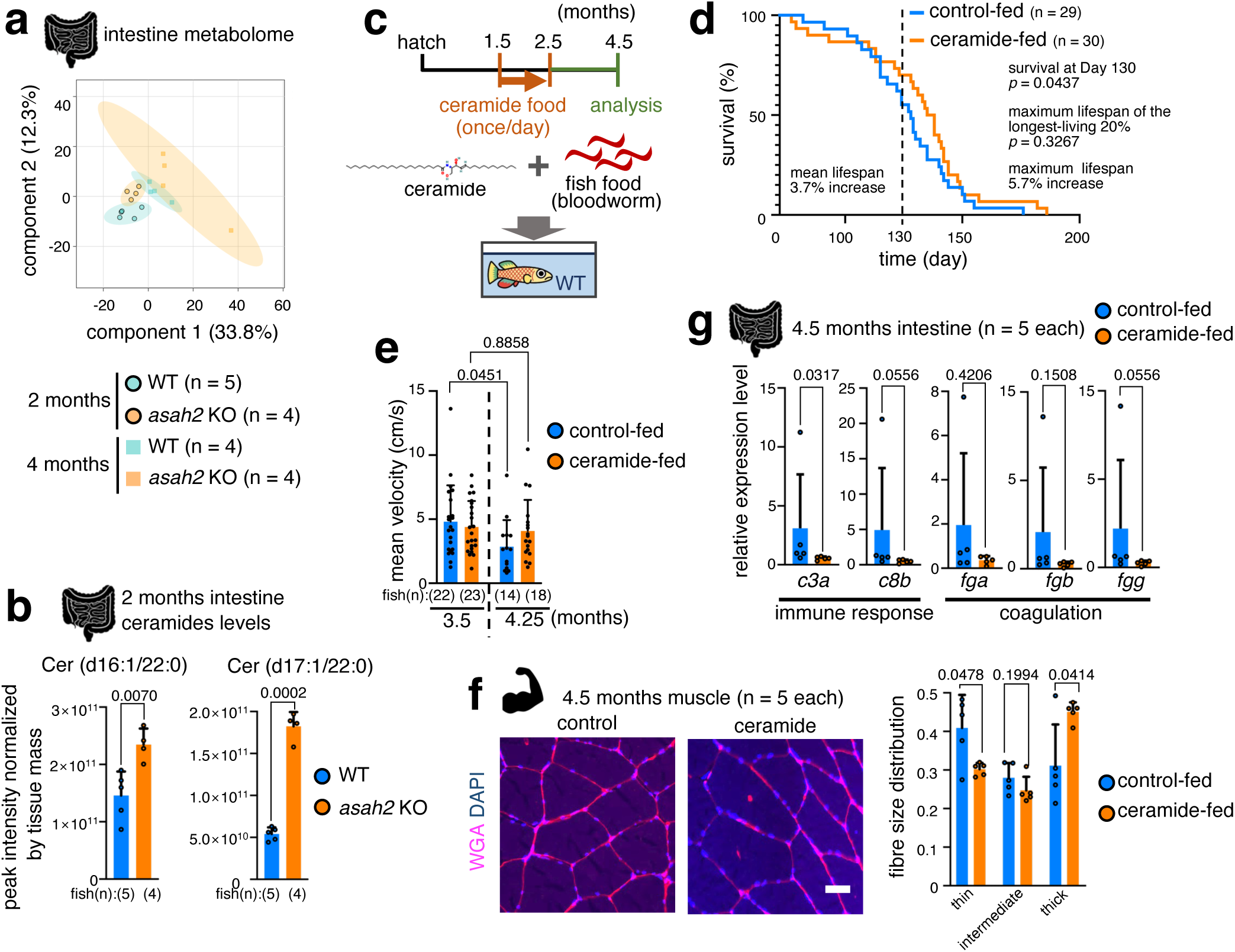
*asah2* KO extends lifespan and healthspan through ceramides. **a**, *asah2* KO alters the intestinal lipidome. Partial least squares–discriminant analysis (PLS-DA) score plot of lipids measured by LC-HRMS-based lipidomics is shown. **b**, *asah2* KO increases Cer(d16:1/22:0) and Cer(d17:1/22:0) in the intestines of 2-month-old fish. The graph shows ceramide levels measured by LC-HRMS-based lipidomics in WT (blue) and *asah2* KO (orange) fish. **c**, Schematic illustration of ceramide supplementation. **d**, **e**, Ceramide supplementation extends lifespan (**d**) and prevents age-related decline in locomotor activity (**e**). Graphs in **d** show survival curves of control-fed (blue) and ceramide-fed (orange) fish. The graph in **e** shows exploratory behaviour in male control-fed (blue) and ceramide-fed (orange) fish. **f, g**, Ceramide supplementation suppresses muscle fibre atrophy and the upregulation of immune response- and coagulation-related genes in aged fish. In **f**, representative images show muscle sections stained with wheat germ agglutinin (WGA; red) and 4′,6-diamidino-2-phenylindole (DAPI; blue). Scale bar, 50 μm. Graphs in **f** show the proportions of thin (< 6,000 μm^2^), intermediate (6,000–9,000 μm^2^), and thick (> 9,000 μm^2^) skeletal muscle fibres in control-fed (blue) and ceramide-fed (orange) fish. In **g**, mRNA levels of immune response- and coagulation-related genes in the intestine of control-fed (blue) and ceramide-fed (orange) fish were analysed by reverse transcription quantitative PCR (RT–qPCR). Bars and error bars (**b, e**–**g**) represent the mean ± SD. An unpaired two-tailed *t*-test was used in **b** and **f**; the log-rank test and fixed-time survival test were used in **d**; a Brown–Forsythe and Welch ANOVA followed by Dunnett’s T3 multiple-comparisons test was used in **e**; and a two-tailed Mann–Whitney U test was used in **g**.

### *asah2* KO suppresses systemic ageing by maintaining microbiota quality

Asah2 is localised in intestinal epithelial cells, and its enzymatic domain is exposed to the extracellular space, facing the intestinal lumen^21^ (Fig. 1a). This suggests that in *asah2* KO fish, ceramides may accumulate extracellularly in the intestinal tract and influence the intestinal environment, including the gut microbiota. Recent studies have shown that gut microbiota regulate systemic aging^16,17,45,46^ and that *asah2* KO alters the microbiota and suppresses liver inflammation and fibrosis in a mouse model of non-alcoholic steatohepatitis^47^. We analysed the gut microbiota using 16S rRNA sequencing (Supplementary Fig. 5a). Interestingly, beta diversity analyses based on Bray-Curtis distances, including principal coordinates analysis and analysis of similarities (ANOSIM), revealed differences in microbial community composition between 2-month-old and 4-month-old wild-type (WT) fish, whereas no such difference was detected between 2-month-old and 4-month-old *asah2* KO fish (Fig. 4a; Supplementary Fig. 5b). In addition, microbiota of ceramide-supplemented fish appeared modestly separated from controls in the PCoA plot (Supplementary Fig. 6a,b). Interestingly, *asah2* KO increased the abundance of *Lactobacillaceae* and *Lactobacillus* at the family and genus levels, respectively (Supplementary Fig. 5a,c). *Lactobacillus* includes probiotic species known to improve the intestinal environment and typically declines with age in humans^48,49^.

**Fig. 4:**
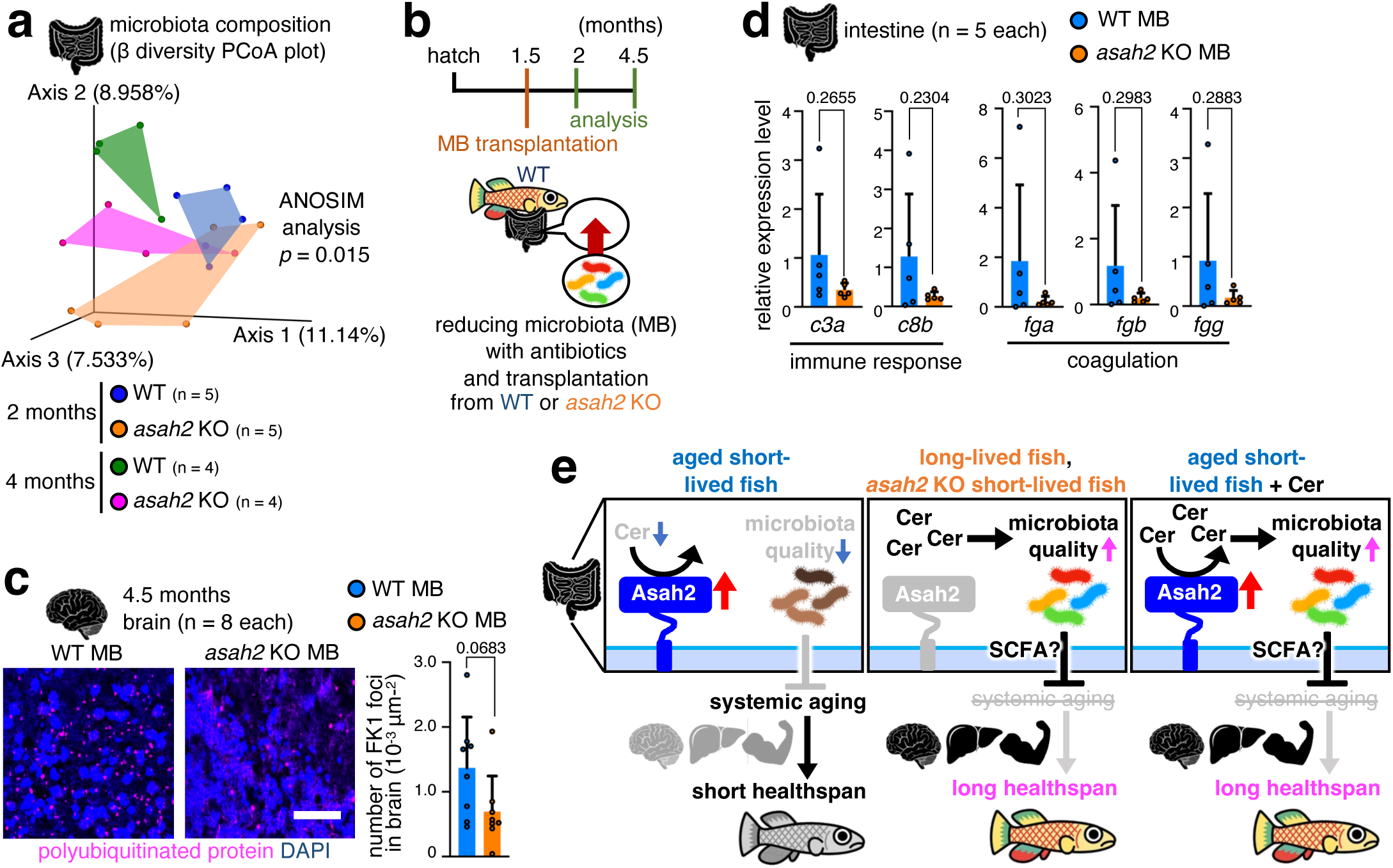
*asah2* KO-derived microbiota prevents ageing. **a**, *asah2* KO attenuates age-dependent changes in microbiota composition. Principal coordinates analysis (PCoA) plots of bacterial composition (β-diversity; a measure of between-sample diversity) based on Bray-Curtis distances are shown. **b**, Schematic illustration of microbiota transplantation. **c, d**, *asah2* KO-derived microbiota suppresses accumulation of polyubiquitinated proteins in the cerebellum and upregulation of immune response- and coagulation-related genes in the intestine of aged fish. In **c**, representative images show cerebellar sections stained with the FK1 antibody (magenta) and 4′,6-diamidino-2-phenylindole (DAPI; blue) in 4.5-month-old fish. Scale bar, 50 μm. The graph in **c** shows the number of FK1 foci in fish transplanted with WT microbiota (WT MB; blue) and *asah2* KO microbiota (*asah2* KO MB; orange) fish. **d**, mRNA levels of immune response-and coagulation-related genes in the intestine of 4.5-month-old fish transplanted with WT MB (blue) and *asah2* KO MB (orange) were analysed by reverse transcription quantitative PCR (RT–qPCR). Bars and error bars (**c, d**) represent the mean ± SD. ANOSIM analysis was used in **a**, and an unpaired two-tailed *t*-test was used in **c** and **d**. **e**, Schematic diagrams illustrating the proposed mechanism by which *asah2* and ceramides regulate the rate of systemic ageing.

*asah2* KO also increased the abundance of *Lachnospiraceae*_NK4A136 and *Muribaculaceae* (Supplementary Fig. 5a,c), which are predicted to produce short-chain fatty acids (SCFAs)^50,51^. *Muribaculaceae* decreases in aged mice^52^, and increases following lifespan-extending interventions, such as caloric restriction and acarbose treatment^53,54^. PICRUSt2-based functional prediction suggested that P124-PWY (*Bifidobacterium shunt*) showed consistently higher median predicted relative abundance in both *asah2* KO and ceramide supplementation (Supplementary Table 3a), a pathway that produces SCFAs, including acetate and ractate^55^. SCFAs suppress age-associated functional decline in the intestine and brain by restoring intestinal barrier and blood–brain barrier integrity, as well as immune homeostasis, and their levels decline with aging^56–58^. Accordingly, the gut microbiota in both *asah2* KO and ceramide-supplemented fish may contribute to prevention of systemic ageing, possibly through production of microbial metabolites. To confirm that *asah2* KO regulates ageing via the microbiota, we transplanted microbiota derived from 1.5-month-old WT or *asah2* KO fish into 1.5-month-old WT recipients (Fig. 4b). This transplantation reduced poly-ubiquitinated protein accumulation in the cerebellum of 4.5-month-old fish (Fig. 4c) and suppressed upregulation of immune response- and coagulation-related genes in 4.5-month-old fish (Fig. 4d). Thus, our findings suggest that *asah2* KO prevents ageing through intestinal ceramide-mediated modulation of the gut microbiota (Fig. 4e).

## Discussion

In this study, we identify the gut-specific ceramidase Asah2 as a contributor to species- and strain-specific determination of ageing rate. Reduced enzymatic activity or protein levels of intestinal Asah2 are associated with extended lifespan in various animals, including *N. furzeri* and humans (Supplementary Fig. 7a). Inhibition of Asah2 and ceramide supplementation alter gut microbiota composition, ameliorate age-related physiological decline, and extend lifespan in the short-lived *N. furzeri* strain (Supplementary Fig. 7b). Our findings suggest that Asah2-mediated ceramide metabolism in the intestine negatively regulates systemic homeostasis and accelerates ageing rates across animal species.

Asah2 is highly conserved among vertebrates^59^, suggesting its indispensability for fundamental cellular and physiological functions. However, Asah2 KO has no apparent adverse effects in mice under normal conditions^23^. Several studies have reported both beneficial and detrimental roles for Asah2 in disease onset and progression. For example, Asah2 KO increases susceptibility of mice to *Citrobacter rodentium* infection^60^, whereas it suppresses liver inflammation and fibrosis^47^. These findings suggest that Asah2 may confer benefits only under specific conditions, such as infection, potentially at the expense of overall health. However, the role of Asah2 in ageing remains unclear. In this study, our comparative genomic analyses predict that several long-lived animals, as well as the long-lived *N. furzeri* strain, lack Asah2 activity. Although we did not experimentally determine whether enzymatic activity of Asah2 is absent in these long-lived species, our findings reveal a previously unrecognised aspect of Asah2 as a lifespan-shortening factor. We also show that, in humans, offspring serum ASAH2 levels are negatively correlated with parental lifespan. The limitations of this analysis are that it assessed the negative correlation between offspring ASAH2 levels and parental longevity, rather than parental ASAH2 levels, and that it measured ASAH2 concentrations in serum rather than levels in the intestine. However, since proteomic profiles are inheritable and Asah2 is expressed predominantly in the human intestine^25–27,30^, it is plausible that intestinal Asah2 also negatively regulates human lifespan.

Notably, in *N. furzeri,* males, but not females, exhibit an age-dependent increase in *asah2* expression (Supplementary Fig. 8a), and *asah2* KO did not extend female lifespan or affect body size or body mass index (Supplementary Fig. 8b–d). In addition, the impact of *asah2* KO on gene expression in the aged intestine of male fish was approximately fivefold greater than that in female fish (Supplementary Fig. 8e). Significant differences in gut microbiota between males and females were also detected, with females showing fewer age-related changes (Supplementary Fig. 8f,g). These findings suggest that Asah2 regulates ageing in a sex-dependent manner, at least in *N. furzeri*. It would be informative to investigate whether Asah2 exerts sex-dependent effects on ageing in other vertebrates, including humans.

Ceramide is a class of sphingolipids, a major group of membrane lipids that constitute the cell membrane. The physiological roles of ceramide have been extensively studied, including its function as an intracellular second messenger, mediator of cell death, and component of the skin barrier^61^. Recent studies have revealed roles for ceramide and related lipid species in regulation of the gut microbiota, tissue homeostasis, and ageing. For example, oral administration of ceramides alters gut microbiota in mice and improves skin conditions in humans^62,63^. Oral administration of glucosylceramide, which can be hydrolysed into ceramide, also alters gut microbiota in mice and improves skin conditions in mice and humans^63,64^. Sphingomyelin, a ceramide precursor, alters gut microbiota in mice^65^, and sphingomyelin- and ceramide-enriched diets ameliorate inflammation in a mouse model of inflammatory bowel disease^63^. Indeed, administration of mixtures of multiple ceramides alters gut microbiota and extends lifespan and healthspan in *C. elegans*^66^. However, the role of ceramides in vertebrate systemic ageing and the specific ceramide species responsible for lifespan extension have remained unclear. In this study, we identify C22 ceramides, Cer(d16:1/22:0) and Cer(d17:1/22:0), as mediators that prevent vertebrate systemic ageing. *asah2* KO increases Cer(d16:1/22:0) and Cer(d17:1/22:0) levels and consequently attenuates intestinal inflammation, increased intestinal permeability (leaky gut), and muscle fibre atrophy. To our knowledge, this is the first study to demonstrate involvement of specific ceramide species in regulation of gut microbiota, systemic ageing, and lifespan in vertebrates. Consistent with our findings, in CerS2 KO mice, decreased C22 ceramide leads to increased intestinal permeability and greater susceptibility to inflammation^67^. These findings suggest that C22 ceramides may also prevent systemic ageing by improving the gut environment in mammals.

Age-dependent decline in gut microbiota quality occurs not only in humans but also in *N. furzeri*^17,46,68^, and such decline is thought to be associated with ageing. Mechanistically, increased gut microbiota diversity and enrichment of SCFA-producing bacteria reduce levels of inflammatory cytokines and strengthen the intestinal barrier^69,70^, whereas gut microbiota dysbiosis (e.g. reduced diversity and increased abundance of pro-inflammatory bacteria) decreases SCFA production and increases intestinal permeability^71^. Indeed, age-related dysbiosis can further increase intestinal permeability and systemic inflammation^16^. In contrast, human centenarians maintain gut microbiota diversity despite advanced age, and preservation of this microbial profile is considered to contribute to healthy ageing^68^. However, few strategies are available to prevent age-associated deterioration of the gut microbiota and to improve healthspan-related phenotypes^46,72^. Our study demonstrates that *asah2* KO and ceramide supplementation improve gut microbiota quality, ameliorate systemic ageing, and extend lifespan. Transplantation of microbiota derived from *asah2* KO fish also improves ageing-related phenotypes in the brain and intestine. These findings suggest that the intestinal Asah2–ceramide–microbiota axis regulates systemic ageing.

Although it remains unclear how the Asah2–ceramide–microbiota axis regulates the brain, muscle, and liver from the gut, one possible mechanism involves modulation of intestinal inflammation. Increased intestinal permeability and inflammation are known to promote systemic inflammation^16,43,44^. Consistent with this interpretation, we show that *asah2* KO, ceramide supplementation, and transplantation of microbiota derived from *asah2* KO fish prevent age-dependent increases in expression of inflammation-related genes, and that *asah2* KO increases expression of Claudin-3, which is required for intestinal barrier function. Another possible mechanism involves gut–brain communication. Recent studies have shown that brain homeostasis is influenced by intestinal inflammation and gut-derived hormones and metabolites^73^. For example, gut dysbiosis and intestinal inflammatory states can activate microbiota–gut–brain/hypothalamic–pituitary–adrenal axis signalling and are associated with elevated systemic inflammatory mediators^74,75^. In intestinal inflammatory conditions, several circulating inflammatory cytokine levels are increased^16,44^, and peripheral challenge with endotoxin or several inflammatory cytokines increases circulating GH levels in humans and sheep^76–78^. Gut microbiota-derived SCFAs also regulate hypothalamic activity^79^, as well as microglial maturation and function in the brain^80^. This study shows that *asah2* KO not only prevents intestinal inflammation and increases abundance of SCFA-producing bacteria in the intestine, but also blocks abnormal protein accumulation and aberrant upregulation of neuroendocrine hormone-related genes, including *gh1* (growth hormone 1) and *prl* (prolactin), in the brain (Supplementary Fig. 3c). These findings suggest that Asah2 regulates brain homeostasis by modulating intestinal inflammation and/or SCFA-producing bacteria (Fig. 4e; Supplementary Fig. 7b). Suppression of abnormal hormone-related gene expression by *asah2* KO in the brain may also contribute to lifespan extension, as inhibition of GH signalling extends mouse lifespan^81^, whereas excess GH shortens lifespan in mice and humans^82,83^, and prolactin is upregulated with age in human males^84^. Notably, GH is known to stimulate mTOR pathway/protein synthesis in peripheral tissues and brain neuronal cells^85,86^. Taken together with our findings that *asah2* KO suppresses age-dependent upregulation of GH expression, increases in ribosomal gene expression, and abnormal protein accumulation in the brain, as well as dysregulated ribosomal gene expression in the liver, *asah2* KO appears to alleviate systemic ageing, likely by suppressing GH upregulation in the brain and the resulting aberrant ribosomal biogenesis and protein synthesis in the brain and other organs. Consistent with this interpretation, elevated ribosome biogenesis is linked to shorter lifespan in *N. furzeri* and is observed in fibroblasts from patients with progeria^37,87^, suggesting that approaches that suppress ribosome biogenesis, such as inhibition of Asah2, could represent an effective strategy for longevity interventions in both humans and fish.

A recent study in *N. furzeri* suggests that ageing can diverge along distinct individual trajectories early in life^37^. However, the upstream factors that determine these divergent ageing trajectories remain unknown. In this context, Asah2 may therefore be viewed not merely as an intestinal lipid hydrolase, but as a potential upstream regulator of divergent ageing trajectories through its effects on the intestinal environment. As ceramides are bioactive lipids that influence the gut environment, altered Asah2 activity may shift the luminal ceramide landscape, thereby reshaping gut microbial composition and function. Such microbiota alterations may, in turn, affect downstream host metabolic signalling and ultimately influence systemic ageing. Although the precise links among ceramides, the microbiota, and systemic regulation of organismal health remain to be fully elucidated, our findings support a model in which intestinal Asah2 modulates organismal ageing by altering a ceramide-centred host–microbiota axis.

## ACKNOWLEDGMENTS

We thank A. Antebi for providing the GRZ strain; B. Hoppe for providing the MZM strain; K. Takeda and R. Okumura, H. Takeuchi and G. Kato for technical advice, and the Ishitani lab members for their helpful discussions, technical support, and fish maintenance. This research was supported by the Takeda Science Foundation (T.I., Y.O.), Nakatani Foundation (T.I.), SECOM Science and Technology Foundation (T.I.), KOSE Cosmetology Foundation (T.I.), Grant-in-Aid for Transformative Research Areas (A) (JP24H02323)(Y.O.) (26H01577)(T.I), Scientific Research (A) (26H02379) (T.I.), Scientific Research (A) (24H02323) (Y.O.), A3 Foresight program (JPJSA3F20230001) (T.I.), MEXT Promotion of Development of a Joint Usage/Research System Project: Coalition of Universities for Research Excellence (CURE) Program (JPMXP1323015484/JPMXP1323015486) (T.I.), Pan-Omics DDRIC, MRCI for High Depth Omics, OU Master Plan Implementation Project promoted under Osaka University (T.I. Y.O.), Young Scientist (20K15701) (K.A.), Kao Foundation for Arts And Sciences (K.A.), AMED-ASPIRE(24jf0126008h0001) (Y.O.) and AMED-CREST (24gm2010001h0001) (T.I.), JST CREST (JPMJCR24T6) (Y.S.), the AMED Moonshot Research and Development Program (JP25zf0127007) (Y.S.), and the AMED Advanced Research and Development Programs for Medical Innovation (JP25gm2010001) (Y.S.), AMED (JP20ek0109492, JP21wm0425009, JP20ek0109485, JP21ek0109548, JP22fk0210111, JP23fk0210138, JP23ek0210194, JP24gm2010001, JP23ek0109675, JP23ek0109672, JP22tm0424222, JP25fk0310536, JP25fk0310535, JP25wm0625519, JP25kk0305031) (M.N.), JST NBDC (JPMJND2302) (M.N.), and JSPS KAKENHI (JP21H02681) (M.N.). This work was partially supported by the “Joint Usage/Research Center for Interdisciplinary Large-scale Information Infrastructures” and the “High Performance Computing Infrastructure” in Japan (Project IDs: jh200047-NWH, jh210018-NWH, jh220014, jh230016, jh240015, and jh250010) (M.N.). This work was supported in part by the MEXT Cooperative Research Project Program for the Medical Institute of Bioregulation, Kyushu University. The infrastructure of the Omics Science Center Secure Information Analysis System (OASIS; https://sis.bioreg.kyushu-u.ac.jp/), Medical Institute of Bioregulation, Kyushu University, provided part of the computational resources. The research has been conducted using the UK Biobank Resource under Application Number 43814 (F.F. and M.N.). F.F. was supported by the Otsuka Toshimi Scholarship Foundation.

## AUTHOR CONTRIBUTIONS

Conception and design: A.T., K.A., and T.I. wrote the main manuscript text; A.T., K.A., and T.I. analysed the data; Y.S. supervised the lipidomic analyses and interpreted the mass spectrometry data; F.F. and M.N. analysed the UK Biobank dataset; A.T., K.A., Y.S., A.K., S.K., H.-N.T., D.M., K.T., M.N., Y.O., and T.I. prepared the figures; A.T., K.A., and T.I. All authors have reviewed the manuscript.

## Methods

### Ethical approval

All experimental animal care was conducted in accordance with institutional and national guidelines and regulations. The study protocol was approved by the Institutional Animal Care and Use Committee of The University of Osaka (RIMD Permit# Biken-AP-R02-04). The study followed the ARRIVE guidelines.

The human data analysis was approved by the Kyushu University Institutional Review Board for Clinical Research (approval number: 23218).

### Fish strain, husbandry, and maintenance

The GRZ (GRZ-AD) strain of *N. furzeri* was kindly provided by A. Antebi (Max Planck Institute for Biology of Aging). The MZM-0410 strain of *N. furzeri* was obtained from B. Hoppe (Leibniz Institute on Aging – Fritz Lipmann Institute). Fish were maintained at 26.5 °C and 0.7 conductivity under a 12/12 h light/dark cycle in a fish breeding system (MEITO, Nagoya, Japan). Adult fish were housed at a density of three per 1.4 L tank from two weeks of age and one per 1.4 L tank from one month of age. Fish were fed freshly hatched brine shrimp twice daily from Monday to Saturday and once daily on Sunday. From two weeks of age, fish were fed bloodworms (Kyorin, Himeji, Japan) once or twice daily on weekdays.

### Variant calling and annotation

High molecular weight DNA was extracted from skeletal muscle using a NucleoBond AXG 500 column (Macherey-Nagel, Düren, Germany). DNA concentration was measured with a Qubit 4 Fluorometer (Thermo Fisher Scientific, Waltham, MA, USA), and size distribution was analysed using a TapeStation 2200 (Agilent Technologies, Santa Clara, CA, USA) to confirm high integrity, with the main peak exceeding 50 kb. A SMRT sequencing library was constructed according to the manufacturer’s protocol using an SMRTbell Express Template Prep Kit 2.0 (Pacific Biosciences, Menlo Park, CA, USA) and sequenced in a single 8M SMRT Cell on a PacBio Revio system (Pacific Biosciences). The output was processed to generate circular consensus sequences, yielding 93.4 Gb of HiFi reads (NCBI BioProject ID 1162561).

Long-read sequencing reads from the MZM strain were aligned to the GRZ genome assembly NfurGRZ-RIMD1 (GCF_043380555.1) using minimap2^1^ (version 2.17-r941). Alignments were sorted and indexed with samtools^2^ (version 1.16.1). Small variants, including single-nucleotide variants and short insertions and deletions, were identified using DeepVariant (version 1.5.0)^3^ using the GRZ assembly as reference. Variants were annotated with snpEff (version 5.3a)^4^, and effects were assigned to overlapping transcripts using Sequence Ontology terms. Coding variants were extracted from the annotated VCF file. Nonsynonymous variants were defined as those annotated with missense_variant, frameshift_variant, stop_gained, start_lost, stop_lost, splice_acceptor_variant, or splice_donor_variant. Variants annotated exclusively as synonymous or non-coding were excluded. Because variants may map to multiple transcript isoforms, annotations were collapsed to the gene level, and each gene was represented once if it harboured at least one nonsynonymous coding variant. When multiple effects were associated with a gene, the most severe predicted effect was retained. Reanalysis using the previous GRZ genome assembly (GCF_001465895.1) and variation and genotype data for MZM-0410 from the *N. furzeri* Information Network Genome Browser (NFINgb)^5^ was performed using snpEff (version 5.3a). All retained variants from both analyses are listed in Supplementary Table 1a, b.

### Characteristic analysis of genes in which SNVs were detected

From the retained SNVs, those located within exons were selected and duplicate transcripts were removed. RefSeq identifiers were converted to Ensembl Gene IDs using db2db in bioDBnet^6^. After removing duplicates, SNVs were identified in 1,669 genes (Supplementary Table 1c).

Overlap between these 1,669 genes and ageing-related gene datasets was examined. Ageing-related disease genes, genes from the GenAge human dataset [Build 21], all genes from the GenAge model organisms dataset [Build 21], and genes associated with longevity variants in humans (LongevityMap, Build 3) were obtained from the Human Ageing Genomic Resources (HAGR)^7^. Killifish Ensembl IDs were converted to gene names using g:Profiler^8^. After excluding non-significant variants in LongevityMap, genes in Supplementary Table 1c were matched against the four datasets using a custom Python script, and overlapping genes were identified (Supplementary Table 1d).

KEGG pathway analysis of the 1,669 genes was performed using ShinyGO (version 0.85)^9^ (Supplementary Table 1e).

To identify metabolic enzymes expressed in the intestine among the 1,669 genes, genes encoding enzymes with Enzyme Commission numbers were retrieved from Ensembl IDs using the UniProt ID mapping tool^10^ (Supplementary Table 2b). Killifish Ensembl IDs were converted to human orthologues using g:Orth in g:Profiler^8^ and Ensembl^11^. Human tissue expression data were obtained from the Human Protein Atlas^12,13^. Metabolic enzymes were curated manually. Tissue-specific metabolic genes are listed in Supplementary Table 2c[NO_PRINTED_FORM].

### Cross-species alignment of *asah2* orthologues

Protein sequences of *asah2* orthologues from multiple animal species, as well as from long- and short-lived *N. furzeri* strains, were aligned using Clustal Omega^14,15^. Protein sequences were retrieved using the following NCBI accession numbers: *Homo sapiens* (NP_063946), *Mus musculus* (NP_061300.1), *Pteropus vampyrus* (XP_011359946.1), *Myotis brandtii* (XP_014402942.1), *Elaphe guttata* (XP_060538933.1), *Perca fluviatilis* (XP_028424295.1), and *Scomber japonicus* (XP_053197810.1).

### Longevity–body mass mapping of Asah2 catalytic-residue-deficient species

The Mammalia, Reptilia, and Teleostei panels were adapted from Kuparinen et al.^16^. Adult body mass and MLS were obtained from the dataset associated with Kuparinen et al.^16^ when available. For catalytic-residue-deficient species not included in Kuparinen et al.^16^, body mass and MLS data were retrieved from AnAge in the HAGR^7^. Body mass and MLS data of these species were ln-transformed and added to the panels as magenta annotations using RStudio^17^ by modifying the code provided by Kuparinen et al.^16^. The original class-averaged regression lines from Kuparinen et al.^16^ were retained. Curated species names, body mass values, MLS values, and data sources are provided in Supplementary Table 4a,b.

### ASAH2 UK Biobank parental longevity analysis

We evaluated the relationship between plasma ASAH2 levels and parental longevity in the UK Biobank using a right-censoring-aware pooled survival framework. ASAH2 was obtained from the UK Biobank RAP Olink normalized protein expression data at instance 0 (olink_instance_0.asah2) and standardized to a z-score. Participant-level covariates were age at the instance-0 assessment (p21003_i0), sex (p31) and genetic principal components 1-20 (p22009_a1–p22009_a20). Genetically inferred European ancestry was defined using p30079 category 5.

Parental longevity was represented by parental attained age. We reconstructed canonical parent attained-age tables from multi-visit UK Biobank source fields for father still-alive status (p1797_i0–p1797_i3), father age at death (p1807_i0–p1807_i3), father current age (p2946_i0–p2946_i3), mother still-alive status (p1835_i0–p1835_i3), mother age at death (p3526_i0–p3526_i3) and mother current age (p1845_i0–p1845_i3). Negative UK Biobank special codes and ages ≤ 18 years were treated as missing, and ages >130 years were discarded. For still-alive status, the latest non-missing response across visits was used. For age-at-death fields, repeated non-missing values were resolved using a predefined mean-if-conflict rule: if conflicting values were present and at least three valid reports were available, their mean was used; otherwise, the latest valid value was retained. For parents alive at the last valid report, current age was defined as the maximum observed current age across visits. Final parental attained age was defined as age at death for deceased parents and current age for living parents, with valid death-age information taking priority over conflicting still-alive or current-age information. Father and mother records were reshaped into long format so that each participant contributed up to two parental rows; living parents were retained as right-censored observations.

The analysis corresponding to Fig. 1f included all eligible participants of genetically inferred European ancestry with complete covariates and at least one parental attained-age record (14,980 participants; 29,412 parental rows; 22,232 events; 7,180 censored observations). We fitted Cox proportional hazards models with parental attained age as the time scale and parental death as the event. Participant plasma ASAH2 z-score was the exposure. Models were stratified by parental sex, defined as mother versus father, to allow different baseline hazards for maternal and paternal rows. Participant-level cluster-robust standard errors were computed using ‘eid’ to account for within-participant dependence between father and mother rows. The primary model was adjusted for participant age, sex and genetic principal components 1–20.

For spline visualization, we fitted cubic B-spline Cox models within the same stratified pooled design using a 3-degree-of-freedom cubic B-spline basis without an intercept. Relative hazards were plotted against ASAH2 z-score with ASAH2 z-score = 0 as the reference. Because likelihood-ratio tests are likelihood-based and do not incorporate the sandwich variance estimator, likelihood-ratio tests for the overall spline association were computed from otherwise identical stratified Cox models fitted without participant-level robust variance; primary inference was based on the cluster-robust linear Cox model.

Sex-stratified follow-up analyses were performed in male and female participants separately. The male follow-up subset contained 6,948 participants contributing 13,621 parental rows (10,460 events; 3,161 censored observations), and the female subset contained 8,032 participants contributing 15,791 parental rows (11,772 events; 4,019 censored observations). These follow-up models used the same pooled long-format parental design and the same parent-type stratification, but were adjusted for measured age and genetic principal components 1–20 only because sex was fixed within each stratum. These sex-stratified analyses were interpreted as lower-powered follow-up analyses rather than primary tests of sex interaction.

### Generation of *asah2* KO *N. furzeri*

A mutant line carrying a 4-bp deletion in *asah2* was generated as previously described^18^. The sgRNA target sites are listed in Supplementary Table 5, and gene sequences were retrieved using the NCBI Gene ID (*asah2*: 107387510). Two target sites within the protein-coding region were selected to avoid overlap with other genomic sequences. Using the CHOPCHOP program, exon 17 was targeted to induce a frameshift mutation and a premature stop codon upstream of the C-terminal region. The sgRNA synthesis protocol followed a previously reported method^18^.

### Genotyping of the *asah2* gene in *N. furzeri*

A 4-bp deletion in *asah2* was detected by PCR using the primers listed in Supplementary Table 5. PCR products were analysed using a microchip electrophoresis system (MCE-202 MultiNA, Shimadzu, Kyoto, Japan) according to the manufacturer’s instructions, with a DNA-500 reagent kit (Shimadzu). The amplified fragments were subsequently sequenced to determine whether they were WT or *asah2* KO.

### Reverse transcription and quantitative polymerase chain reaction

Tissue samples were homogenised in TRIzol reagent (#15596018, Invitrogen, Waltham, MA, USA), and total RNA was extracted using the RNA Clean & Concentrator kit (#R1017, Zymo Research, Irvine, CA, USA). For quantitative PCR (qPCR) analysis, complementary DNA (cDNA) was synthesised using the ReverTra Ace qPCR RT Master Mix with genomic DNA remover (#FSQ-301, Toyobo, Osaka, Japan). qPCR was performed using a Stratagene Mx3000P qPCR system (Agilent Technologies) or CFX Duet Real-Time PCR System (Bio-Rad, Hercules, CA, USA) with THUNDERBIRD SYBR qPCR Mix (#QPS-201; Toyobo) under the following conditions: 95 °C for 1 min, followed by 45 cycles at 95 °C for 15 s and 60 °C for 35 s, or using THUNDERBIRD Next SYBR qPCR Mix (#QPX-201, Toyobo) under the following conditions: 95 °C for 1 min, followed by 45 cycles at 95 °C for 5 s and 60 °C for 35 s. A standard curve method was used to determine relative mRNA abundance. The *tbp* gene was used as a normalisation control. All primers used are listed in Supplementary Table 5.

### RNA-seq: Bulk RNA barcoding and sequencing (BRB-seq)

Tissue samples were homogenised in TRIzol reagent (#15596018, Invitrogen). Total RNA was extracted using the RNA Clean & Concentrator kit (#R1017, Zymo Research), followed by DNA digestion using the DNase I Set (#E1011, Zymo Research). BRB-seq^19^ was performed for library preparation with the following modifications. An oligo(dT)-based primer was used for single-stranded synthesis, and the Second Strand Synthesis Module (#E6111, NEB) was used for double-stranded cDNA synthesis. In-house MEDS-B Tn5 transposase^20,21^ was used for tagmentation, and libraries were amplified by 10 cycles of PCR using Phusion High-Fidelity DNA Polymerase (#M0530, Thermo Scientific, Waltham, MA, USA). Insert reads of 81 bp (Read 2) were generated on an Illumina NovaSeq 6000 platform (Illumina, San Diego, CA, USA).

### RNA-seq: Data processing (BRB-seq)

Read 1 (barcode read) was extracted using UMI-tools (v1.1.1) with the following command: “umi_tools extract -I read1.fastq --read2-in=read2.fastq --bc-pattern=NNNNNNNNNNCCCCCCCCC --read2-stdout”. Adapter and low-quality sequences were removed, and reads shorter than 20 bp were discarded using Trim Galore (version 0.6.10). The processed reads were mapped to the *N. furzeri* Nfu_20140520 reference genome using HISAT2 (version 2.2.1). Read counts for each gene and each sample were obtained by summarising BAM files generated by featureCounts (version 2.0.4). Raw RNA-seq count data were analysed in R (version 4.3.3) with Bioconductor (version 3.18) using edgeR (version 3.42.4). For each sex and tissue, 4-month mutant and wild-type samples were compared separately. Genes with a maximum raw count of fewer than three across the libraries used for filtering were removed before model fitting. Library normalization was performed using the trimmed mean of M-values (TMM) method. Differential expression was assessed using edgeR quasi-likelihood F-tests with legacy = TRUE. P values were adjusted for multiple testing by controlling the false discovery rate (FDR) using the Benjamini–Hochberg method. KEGG pathway and GO term enrichment analyses were performed using ShinyGO 0.85^9^.

### Western blotting

Tissue samples were stored at −80 °C until use. Intestinal specimens were lysed in buffer containing 20 mM Hepes-KOH (pH 7.4), 75 mM NaCl, 12.5 mM β-glycerophosphate, 1.5 mM MgCl_2_, 2 mM EGTA (pH 8.0), 10 mM NaF, 2 mM DTT, 0.5% Triton X-100, 1 mM Na3VO4, 1 mM PMSF, 0.5% aprotinin (014-18113, FUJIFILM Wako Pure Chemical Corporation, Osaka, Japan), protease inhibitor (25955-11, Nacalai Tesque, Kyoto, Japan), and phosphatase inhibitor (07575-51, Nacalai Tesque). Insoluble fractions were removed by centrifugation. Samples were separated by SDS-PAGE, and proteins were transferred to a PVDF membrane (WSE-4051, ATTO, Tokyo, Japan). The membrane was incubated with the appropriate primary antibody: rabbit anti-Asah2 (a custom rabbit polyclonal antibody generated by Eurofins Genomics K.K. (Tokyo, Japan), 1:400), mouse anti-α-tubulin (T6074, Sigma-Aldrich, St. Louis, MO, USA; 1:1000), mouse anti-β-actin (66009-1-Ig, Proteintech, Rosemont, IL, USA; 1:1000), or rabbit anti-claudin-3 (N24P8, Selleck Chemicals, Houston, TX, USA; 1:300), followed by incubation with horseradish peroxidase-conjugated goat anti-rabbit or anti-mouse IgG (401315 and 401215, respectively, EMD Millipore, Burlington, MA, USA). Immunoreactive proteins were visualised using Chemi-Lumi One L (07880-70, Nacalai Tesque) and a FUSION-FX7.EDGE imaging system (M&S Instruments, Osaka, Japan).

### Fertility analysis

Fish fertility was evaluated as previously described with minor modifications^22,23^. Three independent age-matched pairs (one male and one female) of the indicated genotypes were housed in the same tank. Sand trays were added in the morning to allow breeding, and embryos were collected and counted in the evening. Unfertilized eggs die shortly after laying and develop an opaque yolk, enabling clear distinction from fertilized eggs. Results are expressed as the number of eggs per genotype per day.

### Lifespan measurement

Housing conditions were as described above. Fish were fed freshly hatched brine shrimp twice daily from Monday to Saturday and once daily on Sunday. Fish were also fed bloodworms once daily from two weeks to one month of age and once or twice daily thereafter, except during the ceramide supplementation. Mortality was recorded daily from one month of age. In the ceramide supplementation, male mortality was documented daily after treatment. Survival curves were compared using a log-rank test. In *asah2* KO males, lifespan was not significantly changed by the log-rank test, likely because mortality of *asah2* KO males caught up with controls at 4.5 months of age. Therefore, survival rates at a fixed time point (85 days) were compared using the R package “ComparisonSurv”^24^. In the ceramide supplementation, lifespan was also not significantly altered by the log-rank test, likely because mortality of ceramide-supplemented fish caught up with controls at approximately 4 and 5 months of age. Therefore, survival rates at 130 days were compared using the R package “ComparisonSurv”^24^. Lifespan analyses were performed using GraphPad Prism (GraphPad Software, San Diego, CA, USA) with the Kaplan–Meier estimator.

### Histology

Tissue samples were fixed in 4% paraformaldehyde in phosphate-buffered saline (PBS) at 4 °C overnight. Samples were incubated sequentially in 10% and 20% sucrose in PBS until the tissues sank, followed by incubation in 30% sucrose in PBS overnight at 4 °C. Tissues were embedded in Tissue-Tek OCT freezing medium (Sakura Finetek, Tokyo, Japan). Sections (brain: 10 μm; other tissues: 8 μm thick) were prepared using a Thermo Fisher HM525NX cryostat and stored at −80 °C until use. SA-β-gal staining was performed as previously described^25^. Briefly, sections were stained with X-Gal staining solution (7.37 mM citric acid, 25.26 mM sodium phosphate, 5 mM K4[Fe(CN)6]·3H2O, 5 mM K3[Fe(CN)6], 150 mM sodium chloride, 2 mM magnesium chloride, and 1 mg/ml X-Gal in distilled water) for 1–2 h (liver) or 12 h (intestine) at 37 °C and counterstained with Nuclear Fast Red (#NFS500; Scy Tek, Logan, UT, USA). Quantification was performed using ImageJ software.

### Immunohistochemistry analysis

Sections were autoclaved in Target Retrieval Solution, Citrate pH 6 (Dako, Glostrup, Denmark) for 15 min at 105 °C for antigen retrieval. Sections were blocked with 10% FBS, 0.2% Triton X-100, and 1% dimethyl sulfoxide in PBS for 1 h at room temperature. Primary antibodies were anti-Asah2 (produced in this study; 1:100) and anti-multi-ubiquitin monoclonal antibody (mAb; D071-3, MBL, Tokyo, Japan; 1:100). Secondary antibodies were anti-rabbit IgG (#A32733, Invitrogen, 1:500) and anti-mouse IgM (#A-20238; Invitrogen, 1:500). Muscle sections were bleached for 2 h using an LED illuminator (TiYO™, Nepagene, Chiba, Japan). Conjugated wheat germ agglutinin (RL-1022, Vector Laboratories, Newark, CA, USA; 1:300) was used to label muscle membranes, as previously described^26^. Sections were mounted with VECTASHIELD Vibrance Antifade Mounting Medium with DAPI (#H-1800, Vector Laboratories) or ProLong™ Diamond Antifade Mountant with DAPI (P36962, Thermo Fisher, MA, USA). Quantification was performed using ImageJ and AIVIA software (Leica Microsystems, Wetzlar, Germany).

### Lipid analysis

Tissue samples were stored at −80 °C until use. To extract total lipids, frozen tissues were mixed with 100 μl of methanol containing 10 mM ammonium formate. The mixture was homogenised and combined with 100 μl of 1-butanol, followed by centrifugation at 12,700 × g for 15 min at 20 °C. The supernatant was transferred to a 0.2-ml glass insert vial with a Teflon-lined cap for liquid chromatography–electrospray ionisation–mass spectrometry analysis.

For lipidomic analysis, a Q-Exactive Focus Orbitrap mass spectrometer (Thermo Fisher Scientific) was coupled to an HPLC system (Ultimate 3000, Thermo Fisher Scientific). Samples were separated on a Thermo Scientific Accucore C18 column (2.1 × 150 mm, 2.6 μm) using a step gradient of mobile phase A (10 mM ammonium formate in 50% acetonitrile with 0.1% formic acid) and mobile phase B (2 mM ammonium formate in acetonitrile/isopropyl alcohol/water, 10:88:2, v/v/v, with 0.02% formic acid). Gradient ratios were 65:35 (0 min), 40:60 (0–4 min), 15:85 (4–12 min), 0:100 (12–21 min), 0:100 (21–24 min), 65:35 (24–24.1 min), and 100:0 (24.1–28 min) at a flow rate of 0.4 ml min^-^1 and 35 °C. The instrument operated in positive and negative ESI modes, performing a full mass scan (m/z 250–1,100) followed by three rapid data-dependent MS/MS scans at resolutions of 70,000 and 17,500. The automatic gain control target was 1 × 106 ions, with a maximum injection time of 100 ms. Source parameters included spray voltage 3 kV, transfer tube temperature 285 °C, S-Lens 45, heater temperature 370 °C, sheath gas 60, and auxiliary gas 20. Data were analysed using LipidSearch (Mitsui Knowledge Industry, Tokyo, Japan) for annotation of major lipid classes, including ceramides, sphingomyelins, and glycerophospholipids. The endogenous peaks were putatively annotated as Cer(d16:1/22:0) and Cer(d17:1/22:0) based on matching accurate mass, retention time, and MS/MS spectra to those of authentic synthetic standards. For Cer(d17:1/22:0), the long-chain-base assignment was supported by LCB-specific MS/MS fragments, because isobaric or near-isobaric structural isomers such as Cer(d18:1/21:0), Cer(d16:1/23:0), and branched-chain acyl species may otherwise overlap. Search parameters were precursor mass tolerance 3 ppm, product mass tolerance 7 ppm, and m-score threshold 3. Peak areas were normalized to tissue weight and log10-transformed. Pooled QC samples and extraction blanks were included to monitor instrument stability and carryover. The processed peak table was analysed using MetaboAnalyst 6.0^27^.

### Chemical synthesis of Cer(d17:1/22:0) and Cer(d16:1/22:0)

Cer(d17:1/22:0) and Cer(d16:1/22:0) were synthesised by coupling sphingosine (Sph; d17:1) and sphingosine (Sph; d16:1) with the N-hydroxysuccinimide ester of docosanoic acid^28^. Sph(d17:1) and Sph(d16:1) were prepared using a previously reported procedure with minor modifications^29^.

Cer (d17:1/22:0): ^1^H NMR (500 MHz, CDCl_3_) δ 6.24 (d, 1 H, *J* = 7.5 Hz), 5.82–5.75 (m, 1 H), 5.56–5.50 (m, 1 H), 4.34–4.29 (m, 1 H), 3.98–3.88 (m, 2 H), 3.74–3.67 (m, 1 H), 2.78–2.72 (m, 2 H), 2.23 (t, 2 H, *J* = 7.7 Hz), 2.05 (q, 2 H, *J* = 5.3 Hz), 1.68–1.20 (m, 58 H), 0.88 (t, 6 H, *J* = 6.9 Hz); ^13^C NMR (125 MHz, CDCl_3_) δ 174.1, 134.5, 129.0, 74.8, 62.7, 54.7, 37.0, 32.4, 32.1, 29.9, 29.8, 29.8, 29.8, 29.7, 29.6, 29.5, 29.4, 29.4, 29.3, 25.9, 22.8, 14.3; HRMS (ESI) *m/z*: found [M+Na]^+^ 630.5801, C_39_H_77_NO_3_ calcd for [M+Na]^+^ 630.5796.

Cer (d16:1/22:0): ^1^H NMR (500 MHz, CDCl_3_) δ 6.24 (d, 1 H, *J* = 7.5 Hz), 5.83–5.74 (m, 1 H), 5.57–5.50 (m, 1 H), 4.32 (t, 1 H, *J* = 4.8 Hz), 3.96 (dd, 1 H, *J* = 3.8 Hz, J =11.3 Hz), 3.93–3.88 (m, 2 H), 3.70 (dd, 1 H, *J* = 2.9 Hz, *J* = 11.2 Hz), 2.85–2.65 (m, 2 H), 2.23 (t, 2 H, *J* = 7.5 Hz), 2.06 (q, 2 H, *J* = 5.3 Hz), 1.70–1.18 (m, 56 H), 0.88 (t, 6 H, *J* = 6.9 Hz); ^13^C NMR (125 MHz, CDCl_3_) δ 174.0, 134.5, 129.0, 74.9, 62.7, 54.6, 37.0, 32.4, 32.1, 29.9, 29.8, 29.8, 29.8, 29.8, 29.7, 29.6, 29.5, 29.4, 29.4, 29.3, 25.9, 22.8, 14.3; HRMS (ESI) *m/z*: found [M+Na]^+^ 616.5641, C_38_H_75_NO_3_ calcd for [M+Na]^+^ 616.5639.

### Analytical methods for synthesis

^1^H and ^13^C NMR spectra were recorded on an AVANCE III 500 MHz spectrometer (Bruker, Billerica, MA, USA). Chemical shifts in ^1^H NMR spectra are expressed in parts per million (ppm; δ) relative to tetramethylsilane (0.00 ppm). Chemical shifts in ^13^C NMR spectra are referenced to the residual solvent signal (CDCl_3_, 77.16 ppm). Data are reported in the following order: chemical shift, multiplicity (s = singlet, d = doublet, dd = double doublet, t = triplet, q = quartet, m = multiplet), coupling constant(s) in Hz, and integration. High-resolution electrospray ionisation time-of-flight mass spectra (ESI-TOF MS) were acquired using a Bruker micrOTOF mass spectrometer (Bruker Daltonics, Germany).

### Ceramide supplementation

Feeding was performed as previously described, with minor modifications^30^. Cer(d17:1/22:0) and Cer(d16:1/22:0) (0.07 mg each) were added to a single serving of food as follows: a mixed ceramide stock containing Cer(d17:1/22:0) and Cer(d16:1/22:0) at 2 μg/μl each, 4 μg/μl total, was prepared in ethanol and stored at −20 °C. Thirty-five microliters of the stock were added to freeze-dried *Chironomus* larvae (bloodworms; Kyorin) and incubated at 4 °C for approximately 3 h. Subsequently, 5% gelatin was added to the bloodworm/ceramide mixture, mixed, frozen, and stored at −20 °C until use. Control fish were fed food prepared with the same volume of ethanol but without ceramides. Fish received gelatin/Chironomus cubes with or without ceramides for 4 weeks at 1.5–2.5 months of age. Frozen cubes were thawed in water before feeding, and uneaten food was removed. Each fish received one cube daily in the morning. During the ceramide supplementation, fish were additionally fed freshly hatched brine shrimp once daily in the afternoon from Monday to Friday and on Sunday, and twice daily on Saturday. Lifespan, locomotor activity, immunohistochemistry, and gene expression using qPCR were analysed as described above. Microbiome profiles were analysed as described below.

### Microbiome analysis

Intestinal contents were stored at −80 °C until use. DNA extraction was performed using the GENE PREP STAR PI-1200 system (Kurabo, Okayama, Japan). The 27Fmod primer (5′-AGRGTTTGATCMTGGCTCAG-3′) and 338R primer (5′-TGCTGCCTCCCGTAGGAGT-3′) were used to amplify the V1–V2 region of the 16S rRNA gene. A MiSeq Reagent Kit version 2 (500 cycles; Illumina) was used to generate 251 bp paired-end reads on a MiSeq platform (Illumina). The resulting FASTQ files were analysed using QIIME 2^31^ (version 2020.2). After sequence denoising with the DADA2 plugin in QIIME 2, amplicon sequence variants were taxonomically assigned using a V1–V2 region-specific SILVA release 138 99% OTU classifier for the V1–V2 region. Beta-diversity analyses were performed in QIIME 2 using the denoised feature table. Functional potential was inferred from 16S rRNA gene amplicon data using PICRUSt2^32^ implemented through the q2-picrust2 plugin in QIIME 2. For descriptive pathway profiling, PICRUSt2-derived pathway abundance tables were converted to within-sample relative abundances by dividing each pathway abundance by the sum of all predicted pathway abundances in the same sample. Group-wise medians were then calculated for each pathway. Pathways were selected for discussion when the direction of the median difference was concordant across three independent analyses, including two *asah2* KO experiments and one ceramide supplementation.

### Transplantation of fish microbiota

Fish microbiota transplantation was performed as previously described^33^. Briefly, recipient fish were placed in tanks at a density of 10 fish per 9 L of water. Recipient fish were treated overnight with a combination of vancomycin (0.01 g/L), metronidazole (0.5 g/L), neomycin (0.5 g/L), and ampicillin (0.5 g/L). Whole intestines were isolated from donor fish, mechanically disrupted, and added to tanks containing recipient fish at a ratio of one donor intestine per two recipient fish. After incubation overnight in the tank water, fish were returned to the original circulation system. Immunohistochemistry and gene expression analyses by qPCR were performed as described above.

### Statistical analyses

Differences between groups were examined using a two-tailed unpaired Welch’s *t*-test, a two-tailed Mann–Whitney U test, a Brown–Forsythe and Welch ANOVA followed by Dunnett’s T3 multiple-comparisons test, ANOSIM analysis, the log-rank test, and a fixed-time survival test. Statistical analyses were performed using Prism 10 (GraphPad Software, San Diego, CA, USA), Microsoft Excel (Microsoft, Redmond, WA, USA), the R package “ComparisonSurv”^24^, and the quasi-likelihood method in edgeR. Unless otherwise stated, *P* values and false discovery rate values < 0.05 were considered statistically significant. Representative images and plots are shown from a single experiment (Figs. 1c–e, h–i; 2a–g; 3a, b, d–g; 4c, d; and Supplementary Figs. 1a; 2c, d; 3a–c; 4a–c; 5c; 6a, b; 8a–g). Representative images and plots were reproduced in at least two independent experiments (Supplementary Fig. 5a, b).

## Data availability

All data supporting the findings of this study are available within the article, its supplementary information, and the associated source data files. Source data are provided with this paper.

---

**Supplementary Fig. 1:**
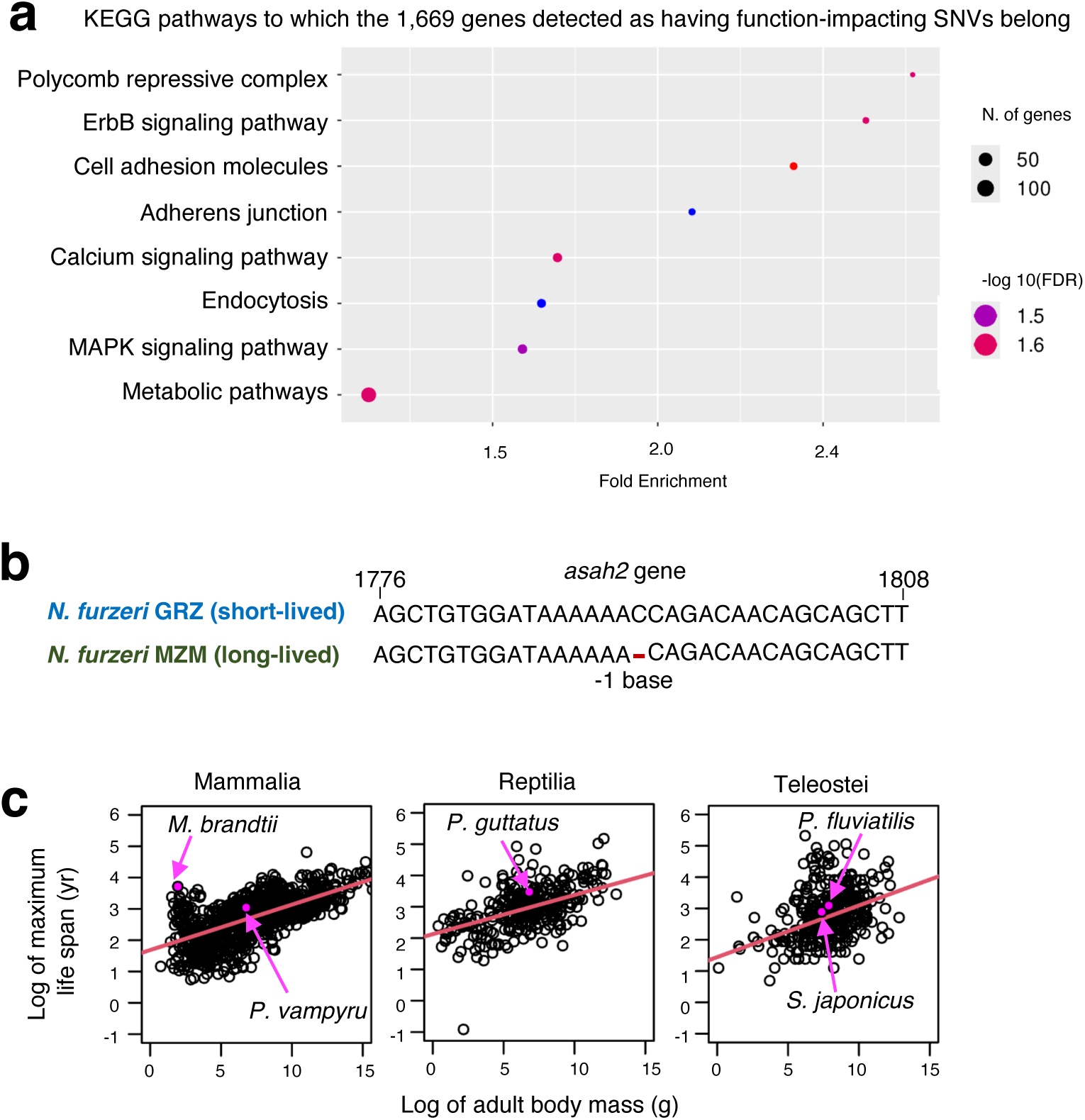
Animals with genetic mutations in Asah2 exhibit longer lifespans. **a**, KEGG pathway analysis of genes harbouring single-nucleotide variants (SNVs) between the short-lived GRZ strain and the long-lived MZM (MZM-0410) strain of *N. furzeri*. **b**, The MZM strain carries a single-nucleotide deletion in the *asah2* gene. The sequences show the nucleotides surrounding the SNV in *asah2* in the GRZ and MZM (MZM-0410) strains. The red horizontal line denotes the deleted base. **c**, Species lacking one or more amino acid residues required for Asah2 enzymatic activity tend to exhibit relatively long lifespans for their body mass within Mammalia, Reptilia, and Teleostei. Log-transformed species-specific average adult body mass and maximum lifespan are shown for these three taxonomic classes. Red lines indicate class-averaged fits from the original linear mixed-effects model. Magenta dots and arrows highlight the corresponding species. Selected panels were adapted from Kuparinen et al.^22^.

**Supplementary Fig. 2:**
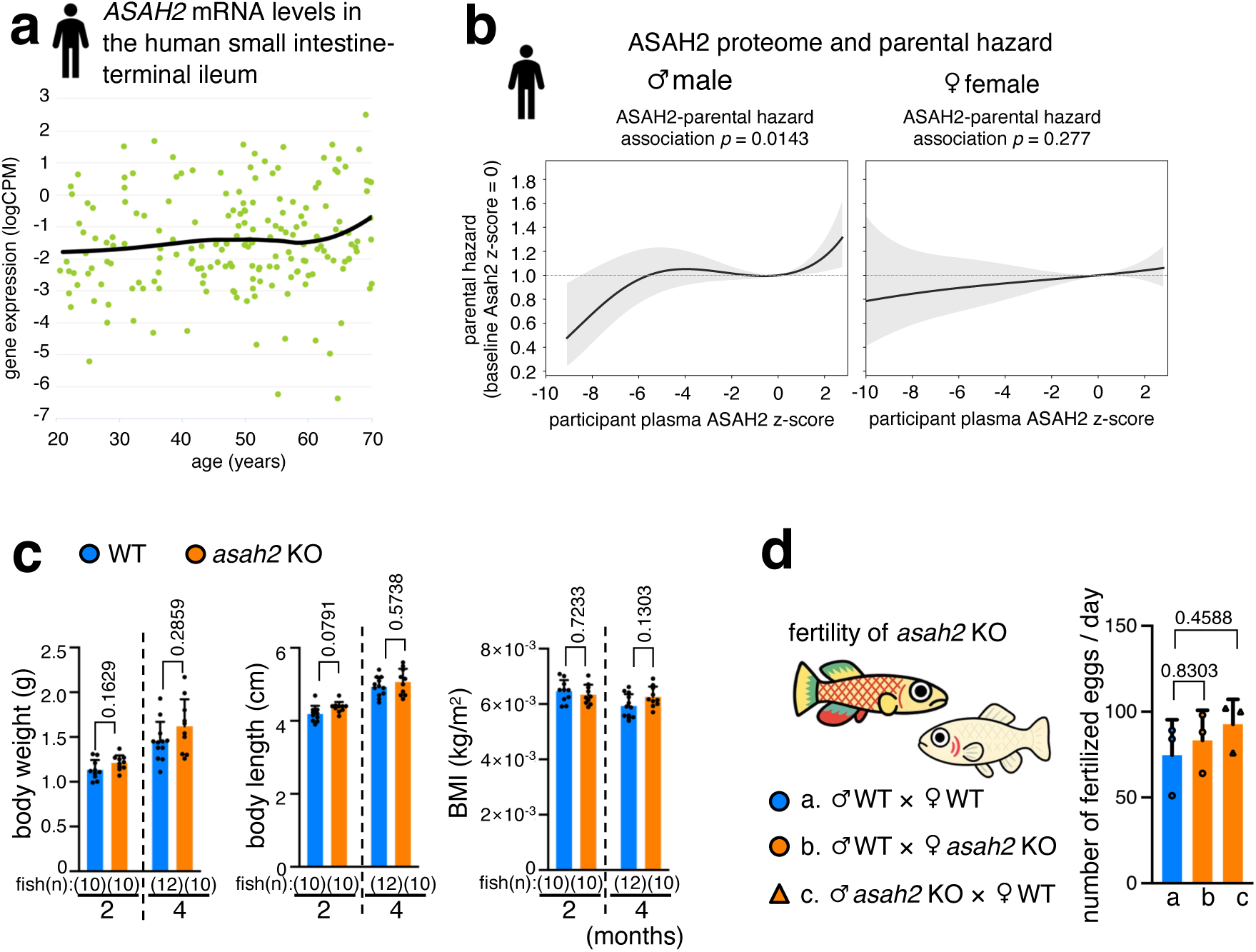
*asah2* mRNA levels increase with age in humans, and *asah2* KO does not affect body size, body mass index, or reproductive capacity. **a**, Asah2 expression increases with age in the human terminal ileum of the small intestine. Data were obtained from voyAGEr^29^. CPM, counts per million. **b**, Sex-stratified pooled censoring-aware parental survival analysis of participant plasma ASAH2 z-score and pooled parental hazard in male and female participants. Displayed p-values denote the overall ASAH2-parental hazard association from stratified cubic B-spline Cox models. **c, d**, *asah2* KO does not affect body size or body mass index (**c**) or reproductive capacity (**d**). Graphs in **c** show body weight, body length, and body mass index in WT (blue) and *asah2* KO (orange) male fish. Animals measured at each time point were different individuals. In **d**, the graph shows the number of fertilized eggs in WT (blue) and *asah2* KO (orange) fish. Bars and error bars (**c, d**) represent the mean ± SD. An unpaired two-tailed *t*-test was used in **c**, and a Brown–Forsythe and Welch ANOVA followed by Dunnett’s T3 multiple-comparisons test were used in **d**.

**Supplementary Fig. 3:**
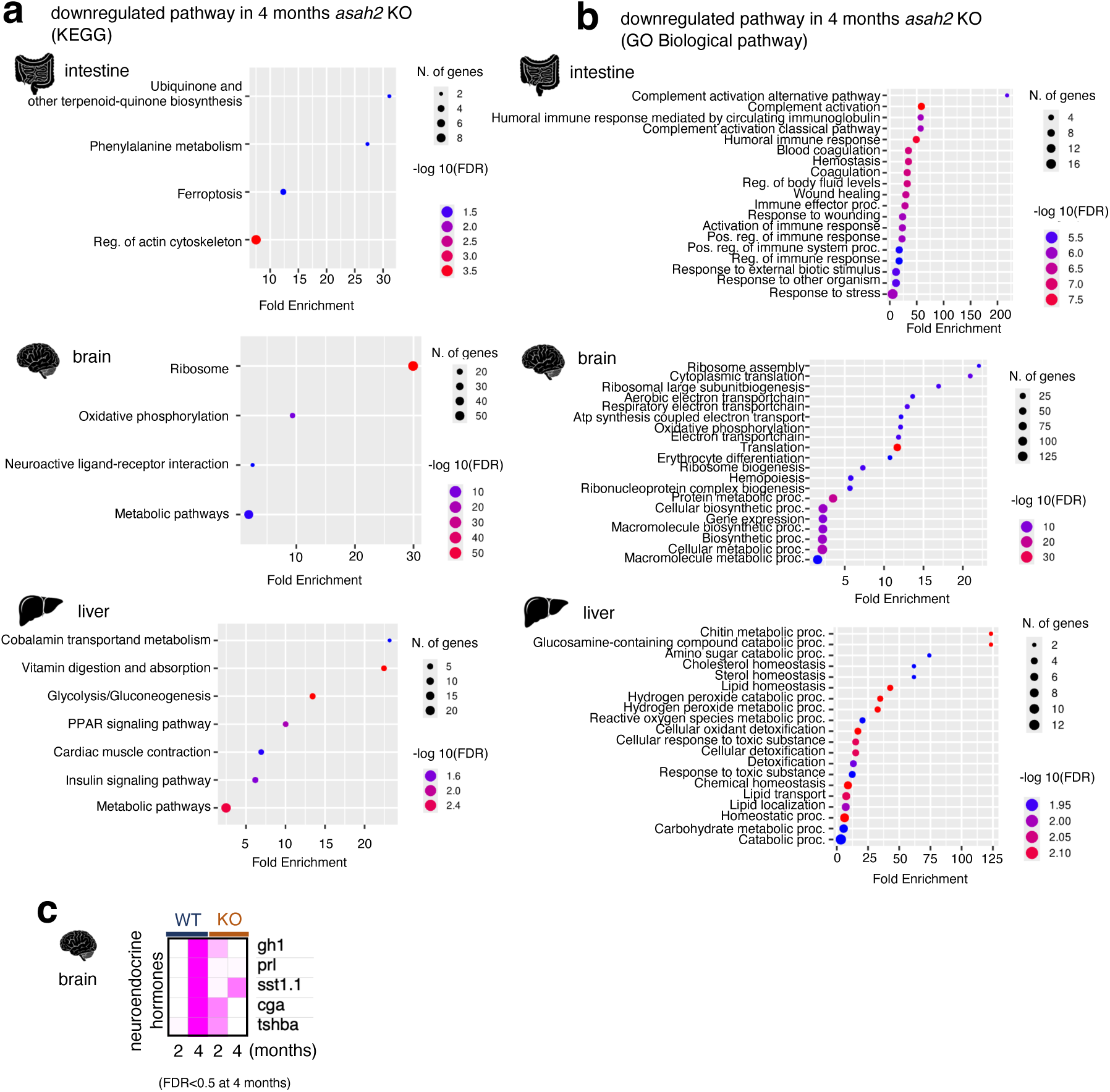
Transcriptomic impact of *asah2* KO. **a, b**, *asah2* KO blocks upregulation of ribosomal protein genes in the brain and complement activation- and blood coagulation-related genes in the intestine of 4-month-old fish. In **a**, KEGG pathway analysis of downregulated genes in 4-month-old *asah2* KO fish is shown. In **b**, Gene Ontology (GO) term enrichment analysis (biological process) of downregulated genes in 4-month-old *asah2* KO fish is shown. For KEGG pathway and GO term enrichment analyses, genes were selected from differential expression analysis using tissue-specific FDR thresholds: FDR < 0.05 for intestine and FDR < 0.5 for brain and liver. **c**, *asah2* KO blocks age-dependent upregulation of neuroendocrine hormone genes in the brain. mRNA levels were analysed by RNA sequencing (RNA-seq). FDR value in **c** was derived from quasi-likelihood methods in edgeR. FDR, false discovery rate.

**Supplementary Fig. 4:**
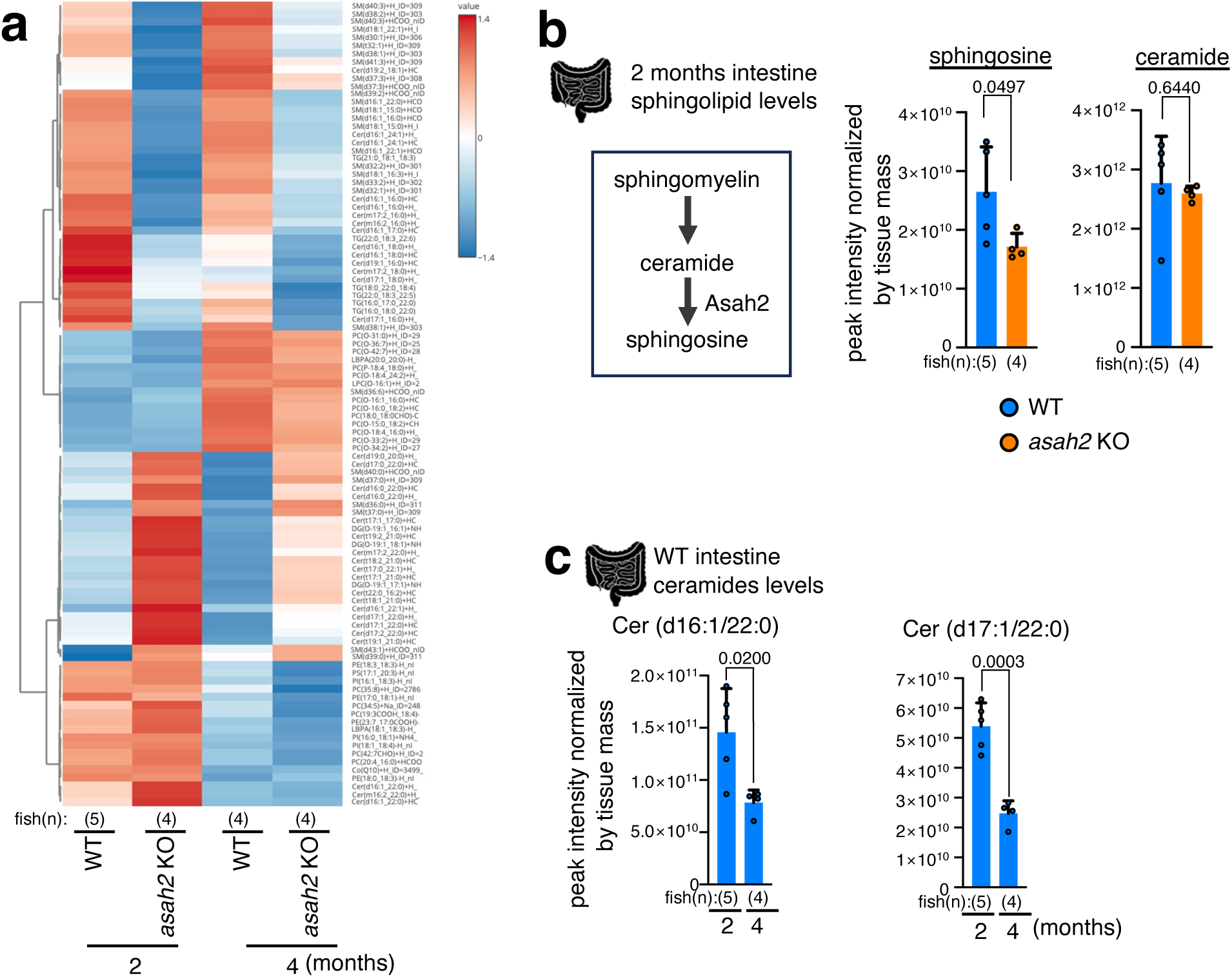
*asah2* KO induces alterations in the lipid profile of male *N. furzeri*. **a**, *asah2* KO alters the intestinal lipid profile. Heatmap of lipids in the intestines of 2- and 4-month-old fish. **b**, *asah2* KO decreases total sphingosine levels but does not change total ceramide levels in the intestine of 2-month-old WT (blue) and *asah2* KO (orange) fish. Levels of total sphingosine and ceramide in the intesine were measured by LC-HRMS-based lipidomics. **c**, Cer(d16:1/22:0) and Cer(d17:1/22:0) decrease with age in WT fish. Bars and error bars (**b, c**) represent the mean ± SD. An unpaired two-tailed *t*-test was used in **b** and **c**.

**Supplementary Fig. 5:**
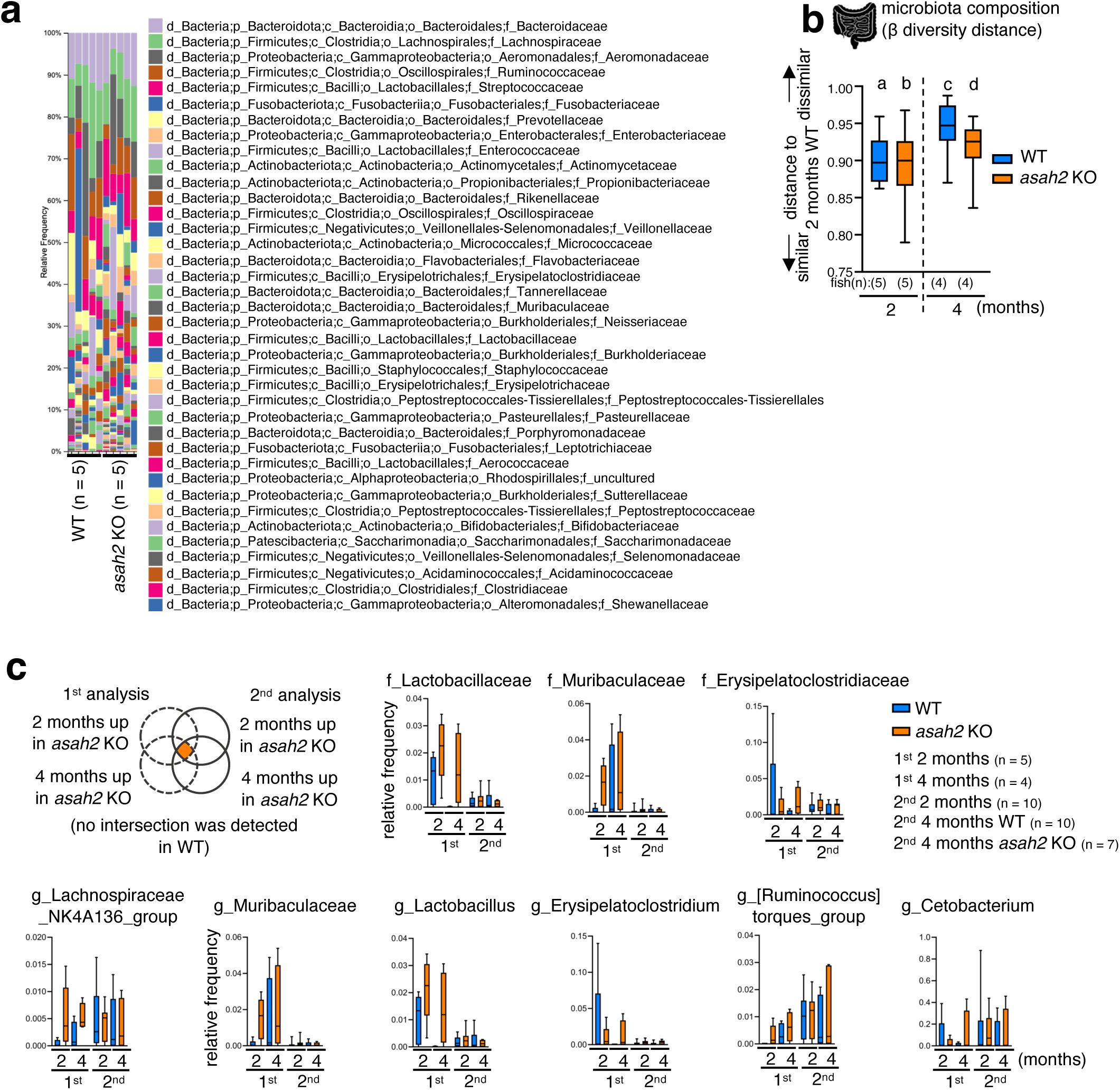
Asah2 regulates microbiota composition. **a**, Relative abundance of the gut microbiota at the family level in WT and *asah2* KO fish. Representative results from two independent experiments are shown. **b**, Box plots of beta diversity (Bray-Curtis distance) and pairwise ANOSIM results for gut microbiota in 2- and 4-month-old WT (blue) and *asah2* KO (orange) fish. Box plots show the distribution of pairwise Bray-Curtis distances. Lower values indicate greater similarity in microbial community composition, whereas higher values indicate greater dissimilarity. For pairwise comparisons, P values were adjusted for multiple testing using the Benjamini–Hochberg method and are reported as q values: a vs b, R = 0.064, q = 0.297; a vs c, R = 0.413, q = 0.093; a vs d, R = 0.213, q = 0.297; b vs c, R = 0.406, q = 0.093; b vs d, R = 0.081, q = 0.297; c vs d, R = 0.094, q = 0.297. **c**, Relative abundance of gut microbial taxa consistently upregulated in *asah2* KO fish. Intestinal microbiota in 2- and 4-month-old WT (blue) and *asah2* KO (orange) fish were analysed. ANOSIM analysis was used in **b**.

**Supplementary Fig. 6:**
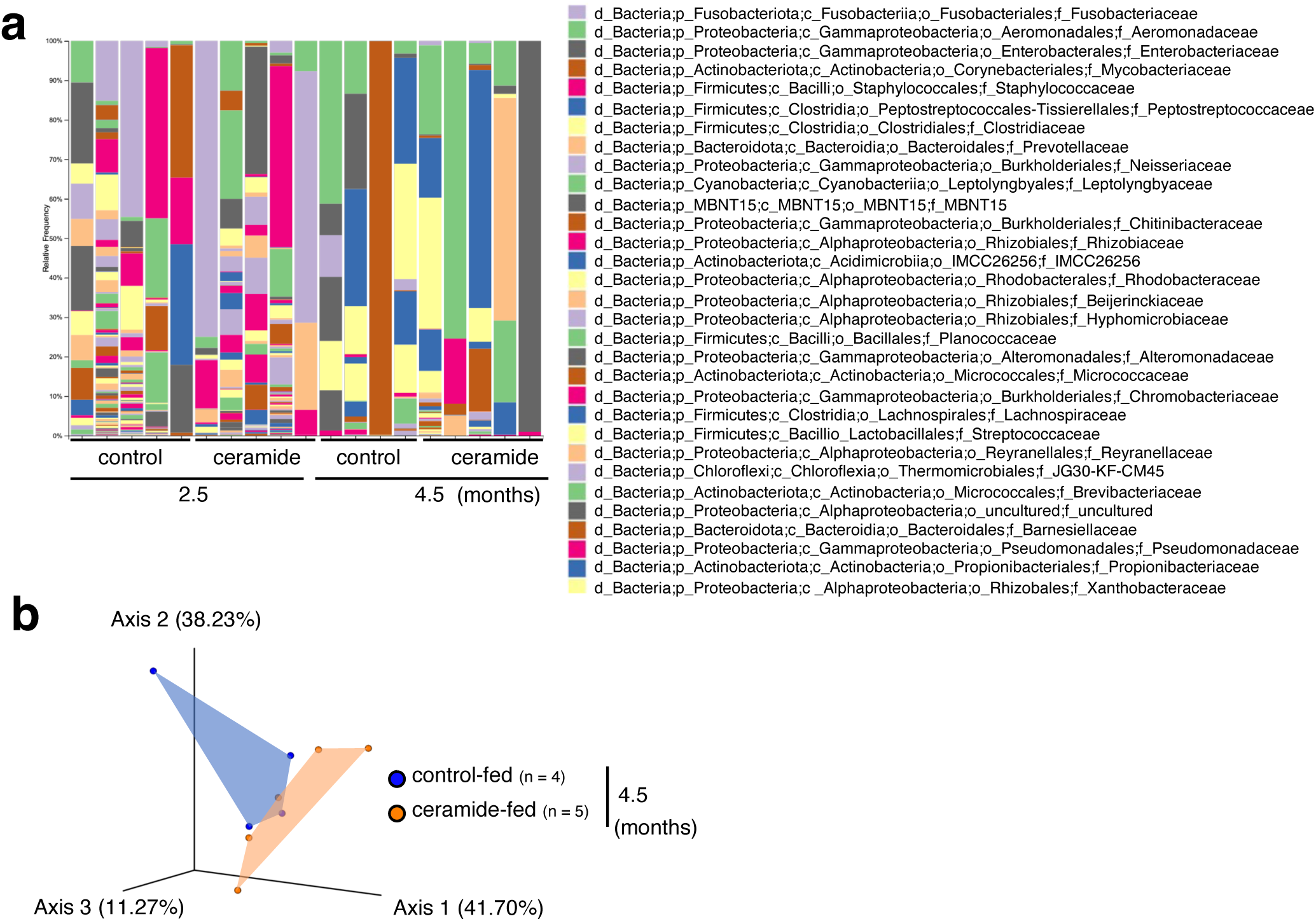
Ceramides alter microbiota composition in male *N. furzeri*. **a**, Relative abundance of the gut microbiota at the family level in control-fed and ceramide-fed fish. **b**, Principal coordinates analysis (PCoA) plots of bacterial β-diversity in control-fed and ceramide-fed fish based on weighted UniFrac distances.

**Supplementary Fig. 7:**
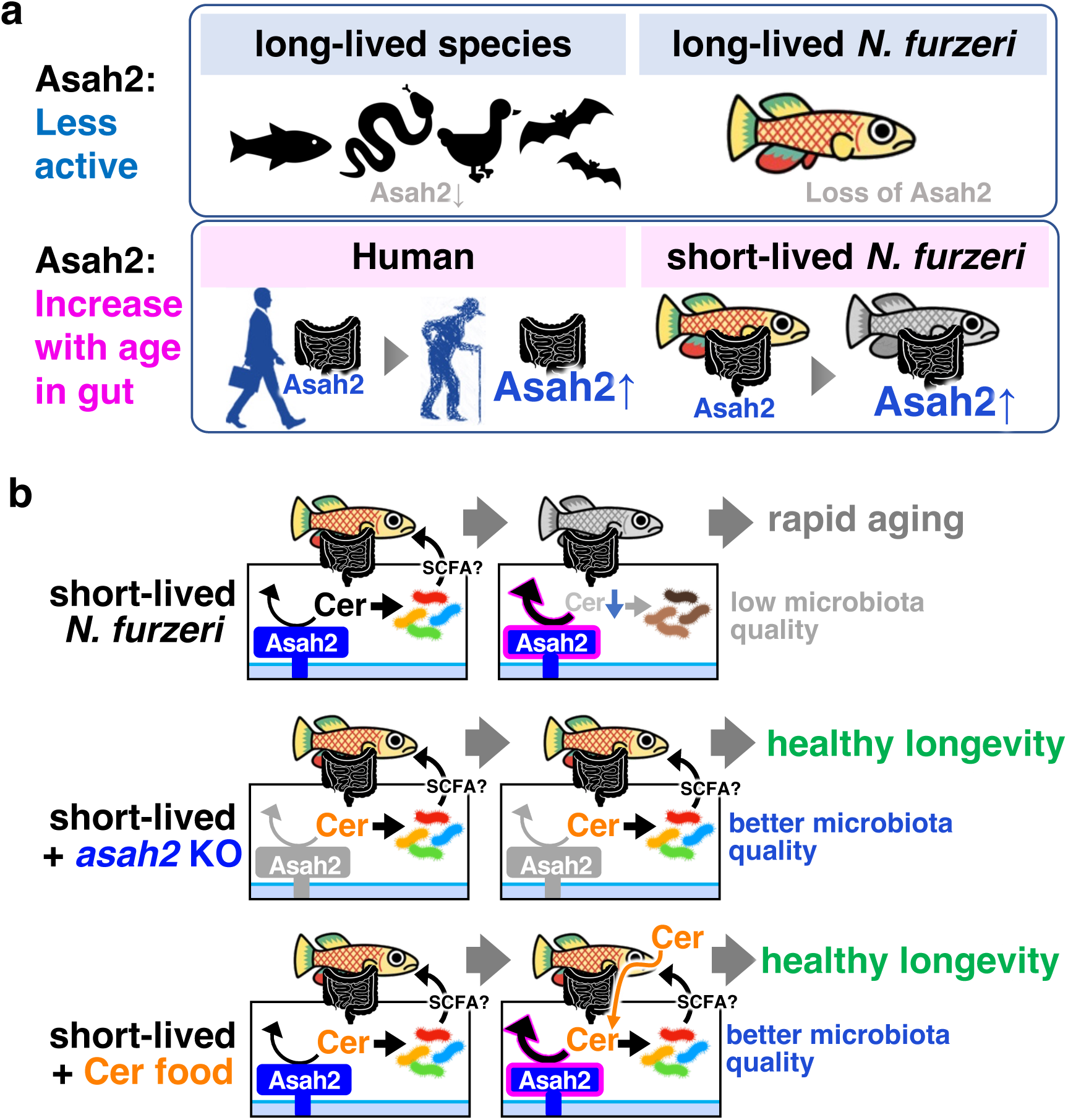
Schematic diagrams illustrating the mechanism by which Asah2 and ceramides regulate systemic ageing through the microbiota. **a**, (Top) Asah2 activity is lower in long-lived species and in the long-lived *N. furzeri* strain. (Bottom) Asah2 expression increases with age in humans and in the short-lived *N. furzer*i strain. **b**, Asah2 accelerates systemic ageing via ceramide-mediated microbiota regulation. *asah2* KO increases ceramide levels, and both *asah2* KO and ceramide supplementation improve microbiota quality and extend healthspan and lifespan.

**Supplementary Fig. 8:**
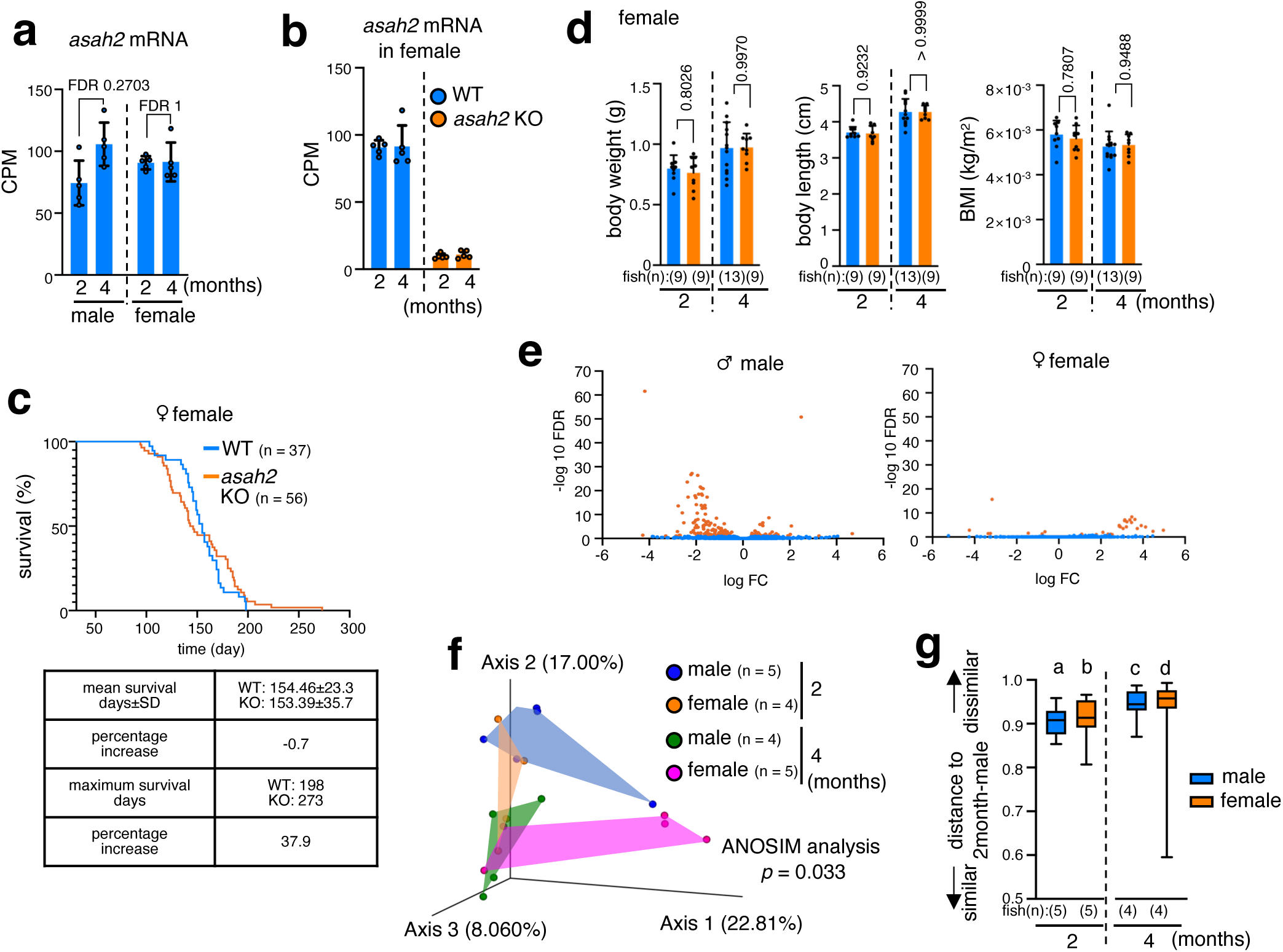
*asah2* KO does not extend lifespan in females. **a**, Expression levels of *asah2* do not change with age in the intestine of female WT fish, as analysed by RNA sequencing (RNA-seq). The data for males are identical to those shown in Fig. 1e. **b**, *asah2* mRNA is depleted in the intestine of female *asah2* KO fish. mRNA levels in WT (blue) and *asah2* KO (orange) fish were analysed by RNA-seq. **c**, Survival curves of female WT (blue) and *asah2* KO (orange) fish are shown at the top; mean lifespan and maximum lifespan are shown at the bottom. **d**, *asah2* KO does not affect body size or body mass index. Graphs show body weight, body length, and body mass index of WT (blue) and *asah2* KO (orange) female fish. Animals measured at each time point were different individuals. **e**, The impact of *asah2* KO on the intestinal transcriptome of 4-month-old fish is greater in males than in female. Volcano plot of differential gene expression in the intestines of 4-month-old WT and *asah2* KO fish. Orange dots indicate genes with FDR < 0.1 (male, 164 genes; female, 30 genes), and blue dots indicate genes with FDR ≥ 0.1. **f, g,** Significant sex differences are observed in the gut microbiota, with females showing fewer age-related changes than males. **f**, Principal coordinates analysis (PCoA) plots of bacterial β-diversity in WT male and female fish based on Bray-Curtis distances. The graph in **g** shows box plots of beta diversity (Bray-Curtis distance) and pairwise ANOSIM results of gut microbiota in 2- and 4-month-old male (blue) and female (orange) WT fish. Box plots show the distribution of pairwise Bray-Curtis distances. Lower values indicate greater similarity in microbial community composition, whereas higher values indicate greater dissimilarity. For pairwise comparisons, P values were adjusted for multiple testing using the Benjamini–Hochberg method and are reported as q values: a vs b, R = 0.088, q = 0.288; a vs c, R = 0.384, q = 0.138; a vs d, R = 0.116, q = 0.268; b vs c, R = 0.115, q = 0.288; b vs d, R = 0.056, q = 0.288; c vs d, R = 0.275, q = 0.268. Bars and error bars (**a, b, d**) represent the mean ± SD. An unpaired two-tailed *t*-test was used in **d**, and ANOSIM analysis was used in **f** and **g**. FDR values in **a** and **e** were derived from quasi-likelihood methods in edgeR. FDR, false discovery rate.

**Supplementary Movie 1**

*asah2* KO ameliorates the age-dependent decline in locomotor activity. Representative movies showing swimming behaviour of young WT fish, aged WT fish and aged *asah2* KO fish are shown on the left, middle and right, respectively. The movie is played at 64× speed.

**Supplementary Table 1**

Single-nucleotide variants (SNVs) predicted to cause loss of protein function between strains and their characteristics. **a,** SNVs predicted to cause a loss of protein function detected in this study. **b,** Single-nucleotide polymorphisms (SNPs) predicted to cause loss of protein function detected using a previous genome assembly and variant dataset. **c,** 1,669 genes with SNVs predicted to cause a loss of protein function. **d,** Genes among the SNV-containing genes that overlapped with known ageing-related genes. **e,** KEGG pathways to which the 1,669 genes detected as having function-impacting SNVs belong.

**Supplementary Table 2**

Filtering steps and identified tissue-specific metabolic enzyme genes among genes containing SNVs. **a,** IDs of 1,669 genes with SNVs predicted to cause loss of protein function, identical to those listed in Supplementary Table 1c. **b,** 339 enzyme-encoding genes with EC numbers among the genes listed in tab a. **c,** Six metabolic enzyme genes with organ-specific expression identified among the genes listed in tab b.

**Supplementary Table 3**

PICRUSt2-based functional analysis of gut microbiota showing the intersection between ceramide feeding and *asah2* KO.

**Supplementary Table 4**

Adult body mass and maximum lifespan data across species. **a,** Species lacking amino acid residues required for Asah2 enzymatic activity and their adult body mass and MLS data. **b,** Original data of adult body mass and MLS. Data from ref. 22.

**Supplementary Table 5**

Primer sequences used in this study.

